# Oncogenic protein condensates suppress growth factor perception and modulate drug tolerance

**DOI:** 10.1101/2022.02.02.478845

**Authors:** David Gonzalez-Martinez, Lee Roth, Thomas R. Mumford, Juan Guan, Bo Huang, Asmin Tulpule, Trever G. Bivona, Lukasz J. Bugaj

**Affiliations:** Department of Bioengineering, University of Pennsylvania, Philadelphia, PA, 19104, USA; Department of Physics, Department of Anatomy and Cell Biology, University of Florida, Gainesville, FL 32611; Department of Anatomy and Cell Biology, University of Florida, Gainesville, FL 32611; Department of Pharmaceutical Chemistry, UCSF, San Francisco CA 94143, USA; Department of Biochemistry and Biophysics, UCSF, San Francisco CA 94143, USA; Chan Zuckerberg Biohub, San Francisco 94158, USA; Division of Pediatric Hematology/Oncology, UCSF, San Francisco, CA 94143, USA; Department of Medicine, Division of Hematology and Oncology, UCSF, San Francisco, CA 94143, USA; Abramson Cancer Center, University of Pennsylvania, Philadelphia, PA, 19104, USA; Institute of Regenerative Medicine, University of Pennsylvania, Philadelphia, PA, 19104, USA

## Abstract

Drug resistance remains a central challenge towards durable cancer therapy, including for cancers driven by the EML4-ALK oncogene. EML4-ALK and related fusion oncogenes form cytoplasmic protein condensates that transmit oncogenic signals through the Ras/Erk pathway. However, whether such condensates play a role in drug response is unclear. Here, we used optogenetics to find that condensates suppress signaling through endogenous RTKs including EGFR. Notably, ALK inhibition hypersensitized RTK signals, which are known to drive resistance. Suppression of RTKs occurred because condensates sequestered downstream adapter proteins that are required for RTK signal transmission. Strikingly, EGFR hypersensitization resulted in rapid and pulsatile Erk signal reactivation, which originated from neighboring apoptotic cells. Paracrine signals promoted survival during ALK inhibition, and blockade of paracrine signals suppressed drug tolerance. Our results uncover a regulatory role for RTK fusion condensates in cancer drug response and demonstrate the potential of optogenetics for uncovering functional biomarkers of cancer cells.

## Introduction

Despite the development of potent oncogene inhibitors, drug resistance remains an unsolved challenge towards durable cancer therapies. Thus, there remains a critical need to better understand molecular interactions between host cells, oncogenes, and targeted inhibitors to identify effective treatment strategies that forestall or prevent resistance.

The oncoprotein EML4-ALK (echinoderm microtubule-associated protein-like 4-anaplastic lymphoma kinase) is a receptor tyrosine kinase (RTK) fusion oncogene that drives ~3-7% of non-small cell lung cancer (NSCLC), the leading cause of cancer-related death in the US (Siegel et al., 2021; Takeuchi et al., 2012). RTK fusions are a large class of chimeric oncogenes that share a similar composition, where the enzymatic fragment of an RTK is fused through chromosomal rearrangement to a fusion partner domain that is often multimeric (Du and Lovly, 2018; Shaw et al., 2013). Cancers driven by RTK fusions commonly exhibit oncogene addiction, where blockade of the oncogene causes cell death(Weinstein, 2002). For EML4-ALK+ cancers, multiple FDA-approved ALK inhibitors achieve initial tumor regression, but resistance and tumor relapse inevitably emerge (Gainor et al., 2016; Solomon et al., 2018).

EML4-ALK drives oncogenicity primarily through sustained Ras/Erk signaling (Hrustanovic et al., 2015). Durable treatment of cancers driven by Ras/Erk has been challenged by robust autoinhibitory feedback loops that result in pathway reactivation after treatment. For example, in the case of BRAF V600E+ cancers, BRAF inhibition suppresses oncogenic Erk signals but simultaneously relieves Erk-dependent negative feedback of RTKs, resulting in strong EGFR stimulation, cell survival, and drug resistance (Corcoran et al., 2012; Gerosa et al., 2020; Lito et al., 2012; Prahallad et al., 2012). For EML4-ALK+ cancers, re-activation of RTK signals after ALK inhibition also promotes drug tolerance and acquired resistance (Hrustanovic et al., 2015; Sasaki et al., 2011; Tani et al., 2016; Vaishnavi et al., 2017; Voena et al., 2013; Wilson et al., 2012) (**Figure 1A**), although specific feedback mechanisms that mediate this reactivation are not well understood.

**Figure 1.**
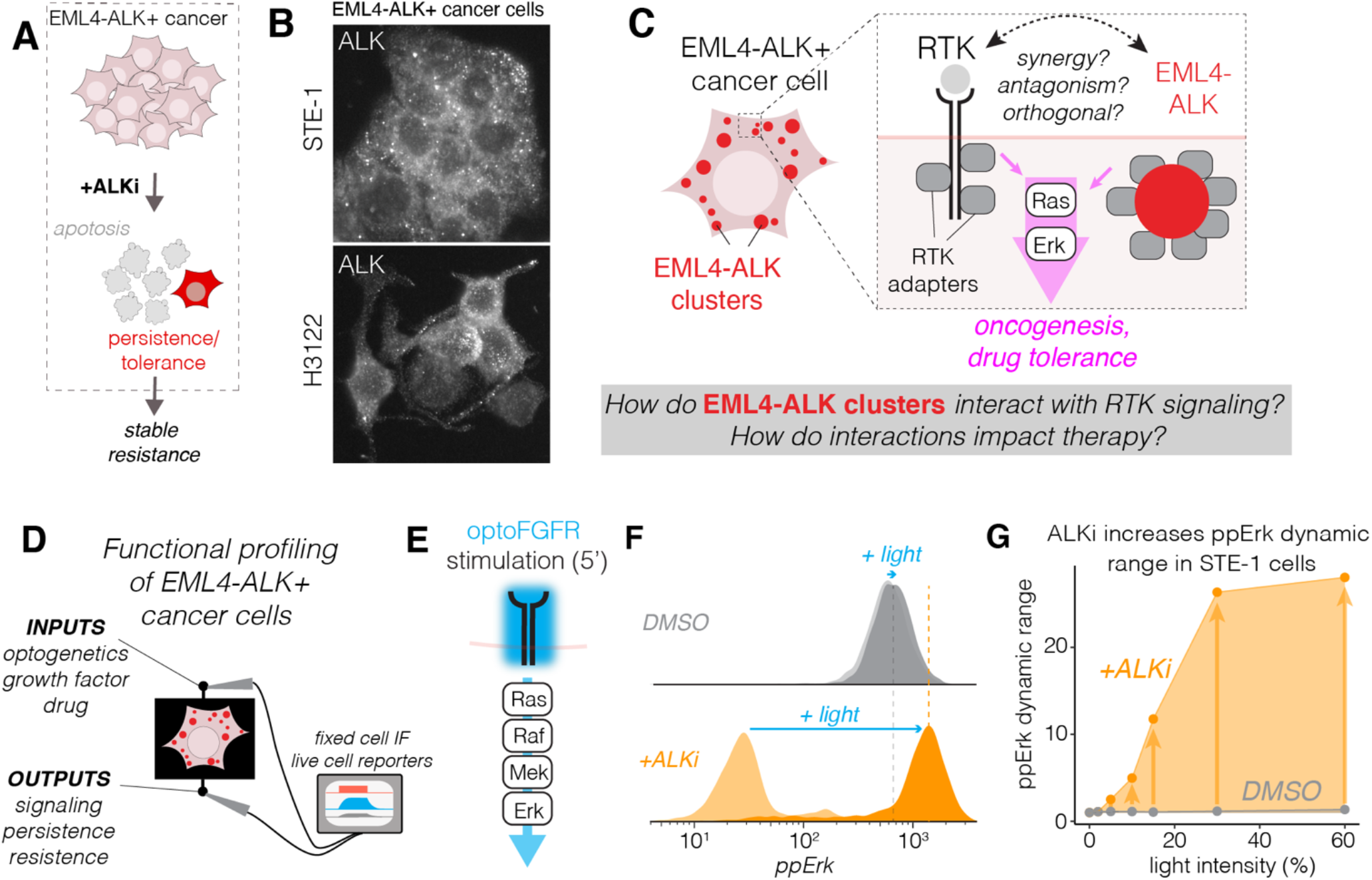
Optogenetics functional profiling of EML4-ALK+ cancer cells reveals suppression of RTK signaling. A) EML4-ALK+ cancer cells treated with ALK inhibitors (ALKi) can persist through therapy and acquire stable drug resistance. B) Immunofluorescence staining shows punctate expression of ALK in two EML4-ALK+ cancer cell lines. C) Although EML4-ALK and transmembrane RTKs signal through the same downstream adapters, the extent to which EML4-ALK condensates interact with RTKs, and whether interactions could impact drug responses, is not clear. D) Functional profiling allows us to observe input/output responses to RTK signaling to understand functional interactions between RTKs and EML4-ALK. E) optoFGFR allows blue-light-induced stimulation of FGFR signaling, including through the Ras/Erk pathway. F) Single-cell immunofluorescence analysis of ppErk levels in STE-1 cells stimulated with light (optoFGFR) in the presence of either DMSO (grey) or the ALK inhibitor crizotinib (ALKi, 1μM; orange). F) ppErk fold-change in response to 5 min of blue light stimulation. Blue light intensity ranged from 2% to 60% of maximum (160 mW/cm^2^). Data points show the ratio of ppErk from stimulated and unstimulated cells. Each condition was performed in duplicate, with each replicate representing the mean of ~4000 cells.

Recently it was found that EML4-ALK forms large cytoplasmic protein granules—or condensates— in cancer cells(Hrustanovic et al., 2015; Tulpule et al., 2021) (**Figure 1B**). Condensate formation resulted in part from the ability of RTK adapter proteins like Grb2 to enlarge small EML4-ALK oligomers through multivalent interactions. Such higher-order assemblies of proteins may have numerous and even opposing consequences on intracellular biochemistry, for example acting as amplifiers or inhibitors of molecular reactions(Banani et al., 2017). Although condensate formation serves as an amplifier that promotes EML4-ALK activation and signaling(Tulpule et al., 2021), it is unclear whether condensates might have additional impact on a cell’s ability to transmit signals. Of particular interest are functional interactions between EML4-ALK condensates and transmembrane RTKs including the epidermal growth factor receptor (EGFR), which signal through the same downstream effectors as EML4-ALK (**Figure 1C**). These interactions are potentially important because EGFR is overexpressed in over 80% of NSCLC and is correlated with poor prognosis (Rusch et al., 1993) and drug resistance (Davies et al., 2013; Lito et al., 2012; Obenauf et al., 2015; Straussman et al., 2012; Tanimoto et al., 2014; Vaishnavi et al., 2017; Wilson et al., 2012).

We recently developed a method called ‘functional profiling’ to detect how oncogene expression can corrupt cell signal transmission (Bugaj et al., 2017, 2018). In this approach, we apply precisely defined signaling stimuli, for example using light-activated ‘optogenetic’ signaling probes, and we quantify differential responses in downstream signals, transcription, and cell fate as a function of oncogene expression or drug treatment. Previously, functional profiling revealed that certain BRAF-mutant cancer cells and drug-treated cells exhibited abnormally slow Ras/Erk pathway activation kinetics, and that such kinetics can cause cells to misinterpret dynamic stimuli leading to hyperproliferation(Bugaj et al., 2018).

In the present work, we applied optogenetics to functionally profile the interaction between EML4-ALK condensates and transmembrane RTK signaling (**Figure 1D**). We discovered that EML4-ALK expression dramatically suppressed a cell’s perception of RTK signals. We found that suppression resulted not from biochemical feedback, but rather from sequestration of RTK adapters by EML4-ALK condensates. Suppression was rapidly reversed by ALK inhibition, resulting in hypersensitization of RTKs to external growth factors. Restored perception of external ligands promoted rapid EGFR/Erk reactivation after drug treatment, which suppressed cell killing and permitted tolerance to ALK inhibitors. Our work thus uncovers an important role for oncogenic condensates in signal regulation and drug response and suggests novel candidate co-targets to enhance the durability of ALK inhibitor therapy.

## Results

### Optogenetic profiling reveals that RTK signaling is suppressed in EML4-ALK+ cancer cells

To determine whether EML4-ALK could alter RTK signal transmission, we expressed a light-sensitive fibroblast growth factor receptor (optoFGFR (Kim et al., 2014)) in the STE-1 cancer cell line, which is driven by EML4-ALK(V1) (Lovly et al., 2014) (**Figure 1E, S1A**). We observed signal transmission by applying blue light stimuli and measuring single-cell phospho-Erk (ppErk) levels through immunofluorescence. We found that optoFGFR could induce only minimal ppErk signal increase (1.4-fold) above the high basal ppErk levels attributed to active EML4-ALK (**Figure 1F, top**). We then asked whether signaling responses might change in the presence of ALK inhibition. Pre-treatment with the ALK inhibitor crizotinib (ALKi, 1 μM) eliminated the high tonic ppErk signal observed in untreated cells, consistent with the ability of EML4-ALK to drive strong signals through the Ras/Erk pathway. Strikingly, subsequent light stimulation now drove strong Erk signaling, surpassing levels achieved in the absence of drug (**Fig. 1F, bottom**). The dynamic range (fold-change) of signal induction increased as a function of light intensity and reached a maximum of 29-fold increase at the highest levels of light stimulation (**Figure 1G**). Notably, ppErk increase was measured in response to exceedingly low levels of light (8 mW/cm^2^), a level that did not provide measurable increase in untreated cells, suggesting that ALK inhibition can hypersensitize cancer cells to weak RTK signals (**Figure S1B**). We note that although the magnitude of dynamic range increase was dependent on the expression levels of optoFGFR, the general trends held across all expression levels (**Figure S1C**). These results suggested the existence of a strong suppressive effect of EML4-ALK activity on transmembrane RTK signaling.

### EML4-ALK suppresses — and ALK inhibition restores — EGFR signaling

To determine whether suppression of RTKs impacted signaling through endogenous epidermal growth factor receptor (EGFR), we measured ppErk induction upon addition of epidermal growth factor (EGF) in STE-1 and H3122 cells, two EML4-ALK+ cell lines (Lovly et al., 2014) (**Figure 2A**). Strong (100 ng/mL) EGF stimulation gave minimal ppErk response, but pre-treatment with ALKi substantially increased ppErk dynamic range in both cell lines (1.8- vs 11.2-fold in STE-; 1.7- vs 6.6-fold in H3122) (**Figure 2B,C**). As before, increased dynamic range was due to lower baseline — but also higher maximal — ppErk signaling, and the magnitude of dynamic range increase was dose-dependent (**Figure 2C, S2A**). Thus, we concluded that EML4-ALK activity suppresses EGFR signaling in cancer cells.

**Figure 2.**
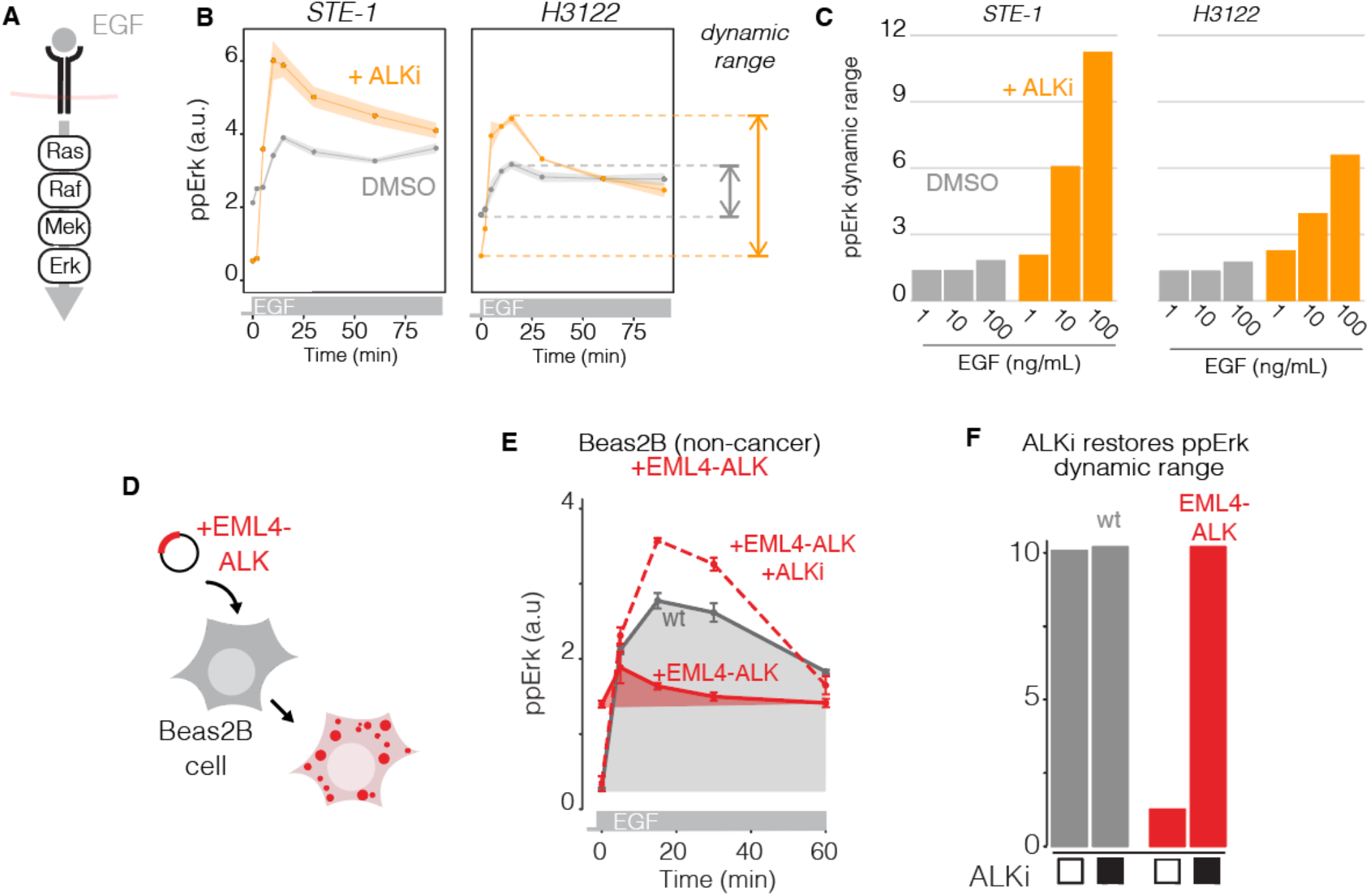
EML4-ALK suppresses EGFR signaling. A) EGF stimulates EGFR and downstream Ras/Erk signaling. B) ppErk levels in response to EGF (100 ng/mL) in the presence of 1μM ALKi (orange) or DMSO (grey) in STE-1 and H3122 cancer cells. Data points and ribbons represent mean +/- SEM of three replicates, each of which represents the mean of ~1000-3000 cells measured through IF. C) Quantification of dynamic range from experiment in 2B over a range of EGF concentrations. D) EML4-ALK(V1) was transiently expressed in lung epithelial Beas2B cells. E) Time course of ppErk immunofluorescence levels in response to EGF stimulation (50 ng/mL). Data points represent mean +/- SEM of three replicates, each of 200-400 transfected or 1000-2000 untransfected cells. F) Dynamic range of ppErk in EML4-ALK-expressing Beas2B in response to EGF in the presence or absence of ALKi (1 μM) pretreatment.

We next asked whether expression of EML4-ALK was sufficient to suppress RTK signaling. We transiently expressed EML4-ALK(V1) in non-transformed lung epithelial Beas2B cells and we observed responses to EGF (**Figure 2D**). EML4-ALK expression raised basal ppErk levels (t = 0, **Figure 2E**), and EGF stimulation resulted in only a small increase in ppErk relative to untransfected cells (**Figure 2E**). Furthermore, pre-incubation with ALKi reversed suppression of EGFR/Erk signals in transfected cells (**Figure 2E,F**), echoing our observations in cancer cells. Collectively, our results show that EML4-ALK expression is sufficient to strongly dampen signaling through transmembrane EGFR, and that ALK inhibition restores and potentiates these signals.

#### Mapping EGFR suppression using optogenetics

We sought to understand the molecular mechanism by which EML4-ALK suppressed EGFR signaling. To narrow candidate mechanisms, we first determined the duration of ALKi pre-incubation that was required to observe enhanced EGFR/Erk signaling. We preincubated both STE-1 and H3122 cells with 0-60 min of ALKi, and we analyzed ppErk levels in response to 15 min of EGF stimulation (**Figure 3A**). In both cell lines, an increase in Erk signal amplitude was observed with as little as 5 min of ALKi pre-incubation (20 minutes total including stimulation), and rose to half-max with only ~15 mins of pre-incubation (**Figure 3B**). Such fast response suggested a primarily post-translational mechanism.

**Figure 3.**
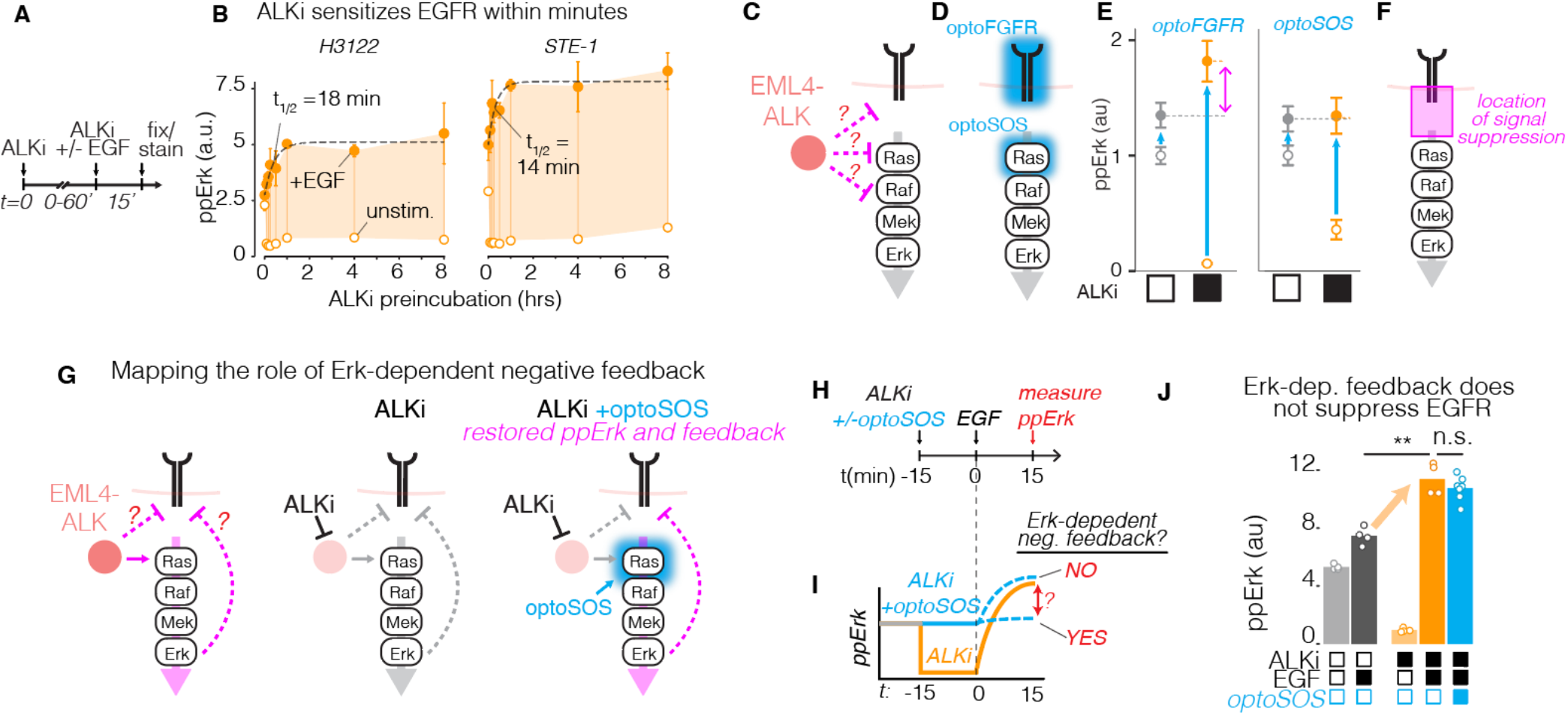
Mapping RTK feedback suppression using optogenetics. A) Time course of RTK hypersensitization was obtained by pre-incubating cancer cells with 1 μM crizotinib for a variable period before stimulation with EGF (50 ng/mL) for 15 min, followed by fixation and immunostaining for ppErk. B) ppErk induction as a function of ALKi pre-incubation time, quantified by immunofluorescence. Open circles = unstimulated. Closed circles = EGF stimulated. Data points represent mean +/- SEM of triplicates, each consisting of 1000-3000 cells. Error bars are smaller than the data points for unstimulated samples. C) Pinpointing the location of EML4-ALK interaction with RTK/Erk signaling. D) OptoFGFR and optoSOS permit optogenetic stimulation at successive nodes of the pathway. E) Quantification of immunofluorescence of ppErk in response to optoFGFR or optoSOS in the presence or absence of ALKi. ALK-dependent suppression is observed only with optoFGFR, suggesting that suppression happens upstream of Ras but downstream of RTK activation (F). G) Testing the role of Erk-dependent negative feedback on RTK suppression. Stimulating optoSOS drives elevated basal levels of ppErk during ALKi treatment and sustains any Erk-dependent negative feedback that would otherwise be lost during ALK inhibition. H) STE-1 cells were treated with either ALKi or ALKi and optoSOS, and EGF response was assessed. I) Predicted results and implications for Erk-dependent feedback. J) Cells treated with both ALKi and optoSOS showed comparable increase in EGF-induced ppErk as cells receiving ALKi only, suggesting that Erk-dependent negative feedback does not suppress EGFR in EML4-ALK+ cancer cells. Data points represent means of ~500-900 cells per condition. ** p < 0.01 by Student’s t-test.

To pinpoint the node in the EGFR/Erk pathway that was suppressed by EML4-ALK, we measured ppErk responses to optogenetic stimulation at successive nodes of the pathway in the presence or absence of drug (**Figure 3C,D**). We stimulated STE-1 cell lines that stably expressed either optoFGFR or optoSOS, which allows activation of Ras/Erk signals through light-induced membrane recruitment of the SOS catalytic domain(Benman et al., 2022; Toettcher et al., 2013) (**Figure 3D, S2**). As before, light stimulation of optoFGFR yielded little increase in ppErk, but ALKi pretreatment enhanced ppErk induction (**Figure 3E, left**), consistent with relief of optoFGFR suppression by ALKi. By contrast, the maximal level of ppErk induced by optoSOS was unchanged by ALKi pre-treatment (**Figure 3E, right**). These results suggest that EML4-ALK suppresses signaling upstream of Ras stimulation, for example at the receptor level. However, further experiments showed that EGF-stimulated phospho-EGFR (pEGFR) levels were unchanged by ALKi pre-incubation, demonstrating that EML4-ALK does not directly suppress RTK phosphorylation (**Figure S3**). Collectively, our results indicate that EML4-ALK suppresses RTK signaling downstream of receptor phosphorylation but upstream of Ras activation, implicating a role for the adapter proteins that couple these two nodes (**Figure 3F**).

Oncogene-mediated feedback suppression of RTKs has been described previously, most notably as a result of Erk-dependent transcriptional and post-translational negative feedback(Corcoran et al., 2012; Gerosa et al., 2020; Lito et al., 2012; Prahallad et al., 2012; Turke et al., 2012). To test the role of Erk-dependent negative feedback in EML4-ALK+ cancer cells, we decoupled ALK inhibition from the loss of Erk signaling using optoSOS stimulation (**Figure 3G**). In this experiment, we again assessed EGF response after ALKi preincubation, but we supplemented one experimental group with optogenetic Ras/Erk signaling at levels that matched those from drug-naïve STE-1 cells, thus maintaining any Erk-dependent feedback that would have been otherwise lost through ALK inhibition (**Figure 3G,H,I, S4A**). We found that optoSOS stimulation during ALKi pre-incubation did not diminish the enhanced response to EGF (**Figure 3J, S4B**). In agreement, the levels of Erk-dependent negative regulator Spry2 did not change within the first 60 min of ALKi pre-incubation (**Figure S4C**). These results show that, in EML4-ALK+ cancer cells, EGFR suppression is not mediated by established, Erk-dependent mechanisms.

#### EML4-ALK condensates suppress EGFR through sequestration of RTK effectors

We suspected that condensation of EML4-ALK could play a causal role in RTK suppression. To test the effects of such higher-order EML4-ALK organization, we examined EGF response in cells transfected with EML4-ALK mutants that fail to form condensates due to mutations in either the trimerization domain (ΔTD) or the kinase domain (K589M)(Tulpule et al., 2021). In contrast to full-length EML4-ALK (V1) (**Figure 4A, left**), the condensate-deficient mutants had minimal effect on EGF-induced Erk activity (**Figure 4A, middle and right**), suggesting that condensation was essential for its ability to modulate EGFR sensitivity.

**Figure 4.**
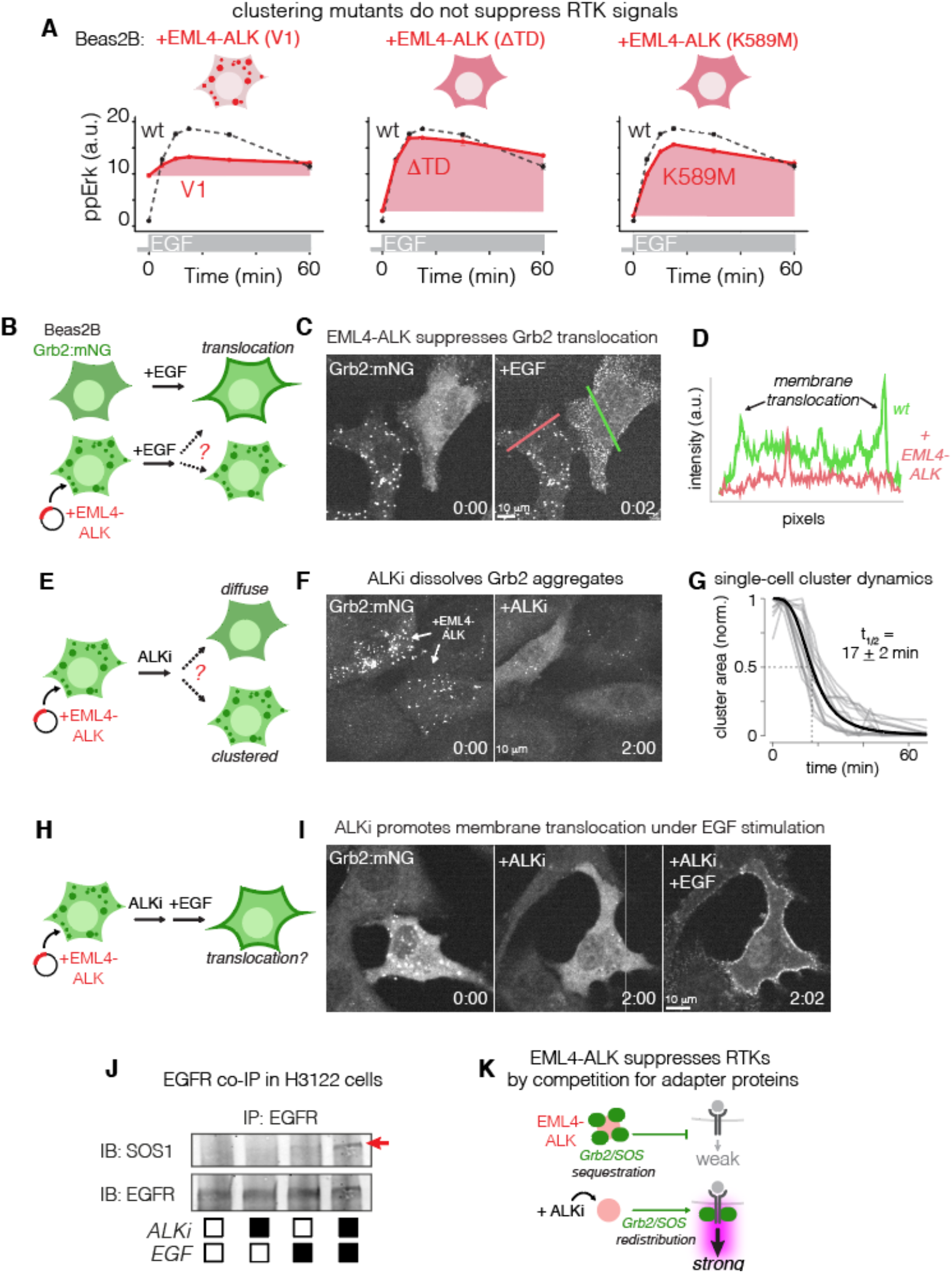
EML4-ALK condensates suppress RTK signals through sequestration of RTK effectors. A) Quantification of ppErk response to EGF (50 ng/mL) stimulation in Beas2B cells that expressed EML4-ALK (V1), EML4-ALK (ΔTD), and kinase-dead EML4-ALK (K589M). Data are means +/- SEM of triplicates, each of which represents 500-1000 cells. Data points represent mean. B) Beas2B cells that harbored endogenously-tagged Grb2 (Grb2:mNG) were transiently transfected with EML4-ALK and stimulated with EGF (50ng/mL) to visualize Grb2 translocation in the presence and absence of EML4-ALK. C) Impaired membrane translocation of Grb2 in the presence of EML4-ALK condensates. Time in mm:sec. See **Supplementary Movie 1**. D) Line scan of Grb2 intensity distribution in the presence (red) or absence (green) of EML4-ALK expression, as depicted in C. E) Grb2 localization was visualized upon treatment with 1 μM ALKi. F) Grb2 shifted from condensates to cytoplasm after ALKi treatment. Time in hh:mm. See **Supplementary Movie 2**. G) Quantification of Grb2 cluster dissociation kinetics after ALKi treatment. H) Grb2 distribution was examined upon sequential treatment with ALKi and EGF. I) ALKi released Grb2 to the cytoplasm, which allowed its membrane translocation in the presence of EGF. J) Immunoprecipitation of EGFR shows co-precipitation of SOS1 only in the presence of ALKi pretreatment and EGF. K) Conceptual model of how EML4-ALK suppresses transmembrane RTKs. EML4-ALK sequesters adapters like Grb2/SOS1 and prohibits their translocation to activated RTKs. ALK inhibition releases adapter sequestration and restores cellular response to RTK stimulation.

Protein condensation can act as a negative regulator in both natural and engineered systems when it sequesters essential components of biochemical reactions (Garabedian et al., 2021; Omer et al., 2018; Thedieck et al., 2013). We thus hypothesized that EML4-ALK condensates suppressed transmembrane RTKs by sequestering the adapter proteins that are required to transmit RTK signals. Such adapters, including Grb2 and SOS1 which link RTKs to Ras/Erk signaling, are also required to transmit EML4-ALK signals (Tulpule et al., 2021), and thus represent shared resources that could implement competitive inhibition of RTKs.

To test this hypothesis, we observed the localization of the Grb2 adapter using Beas2B cells where Grb2 was fluorescently tagged at the endogenous locus (Tulpule et al., 2021). In normal cells, Grb2 appeared diffuse in the cytoplasm but translocated to activated EGFR within <1 min of EGF addition (**Fig. 4B,C, Supplementary Movie 1**). However, upon transfection with EML4-ALK, Grb2 strongly colocalized with EML4-ALK puncta (**Figure S5**), and Grb2 remained sequestered in the puncta even upon addition of EGF (**Figure 4B,C,D, Supplementary Movie 1**). We then asked how Grb2 localization might change in the presence of ALKi (**Figure 4E**). ALKi caused Grb2 puncta to rapidly dissolve into the cytoplasm (t_1/2_ = 17 +/- 2 min) (**Figure 4F,G, Supplementary Movie 2**), in line with recent reports (Sampson et al., 2021). Finally, sequential treatment with ALKi and EGF allowed robust membrane translocation of Grb2 even in the presence of EML4-ALK (**Figure 4H,I**). These results suggest that EML4-ALK condensates sequester Grb2 and suppress translocation to the membrane upon EGF stimulation, providing a mechanism by which EML4-ALK can competitively inhibit EGFR signaling.

As a further test of our model, we directly measured the EGF-induced recruitment of adapters in H3122 cancer cells through co-immunoprecipitation. SOS1 is recruited by Grb2 to activated receptors and stimulates Ras upon recruitment. We found that SOS1 co-precipitated with EGFR upon EGF stimulation, but only when preincubated with ALKi, further suggesting that active EML4-ALK condensates sequester RTK adapters (**Figure 4J, S6**). Finally, our conceptual model predicts that distinct RTK fusions that form condensates would similarly repress EGFR signaling. We thus measured EGF response in TPC-1 cells, which harbor the CCDC6-RET fusion. Like EML4-ALK, CCDC6-RET forms condensates that colocalize with many RTK adapters, including Grb2 and SOS1(Tulpule et al., 2021). As predicted, EGF stimulation induced only moderate levels of Erk phosphorylation, whereas pre-treatment with RET inhibitor BLU-667 permitted strong stimulation (2.0- vs 6.0-fold stimulation) to levels beyond those achievable in drug-naive cells (**Figure S7**). Collectively, our results suggest that competition for RTK adapters is a key mechanism by which condensates of EML4-ALK —and potentially other RTK fusions — suppress a cell’s perception of transmembrane RTK signals (**Figure 4K**).

### RTK resensitization promotes rapid signal reactivation upon ALK inhibition

RTK stimulation promotes drug tolerance and acquired resistance across cancer types (Straussman et al., 2012; Wilson et al., 2012). We thus sought to understand whether hypersensitization of RTKs could promote RTK signaling during ALKi therapy. We monitored RTK/Erk signaling in drug-treated populations of STE-1 cells using ErkKTR, a biosensor that reports on Erk activity through nuclear exclusion of a fluorescent protein (Regot et al., 2014). (**Figure 5A**). In the absence of ALKi, Erk activity was at an intermediate level, indicated by relatively equal distribution between the cytoplasm and nucleus (**Figure 5B, Supplementary Movie 3**). Notably, this localization pattern did not change over the course of 22 hrs, reflecting the tonic signaling downstream of EML4-ALK. By contrast, treatment with ALKi induced a rapid initial decrease of Erk activity followed by a striking appearance of Erk activity pulses within ~1 hr after treatment (**Supplementary Movie 3**). Each Erk pulse lasted 10-20 minutes, and the pulse amplitude exceeded levels observed in the absence of drug, paralleling our earlier results with optogenetic or ligand stimulation (**Figure 5C**).

**Figure 5.**
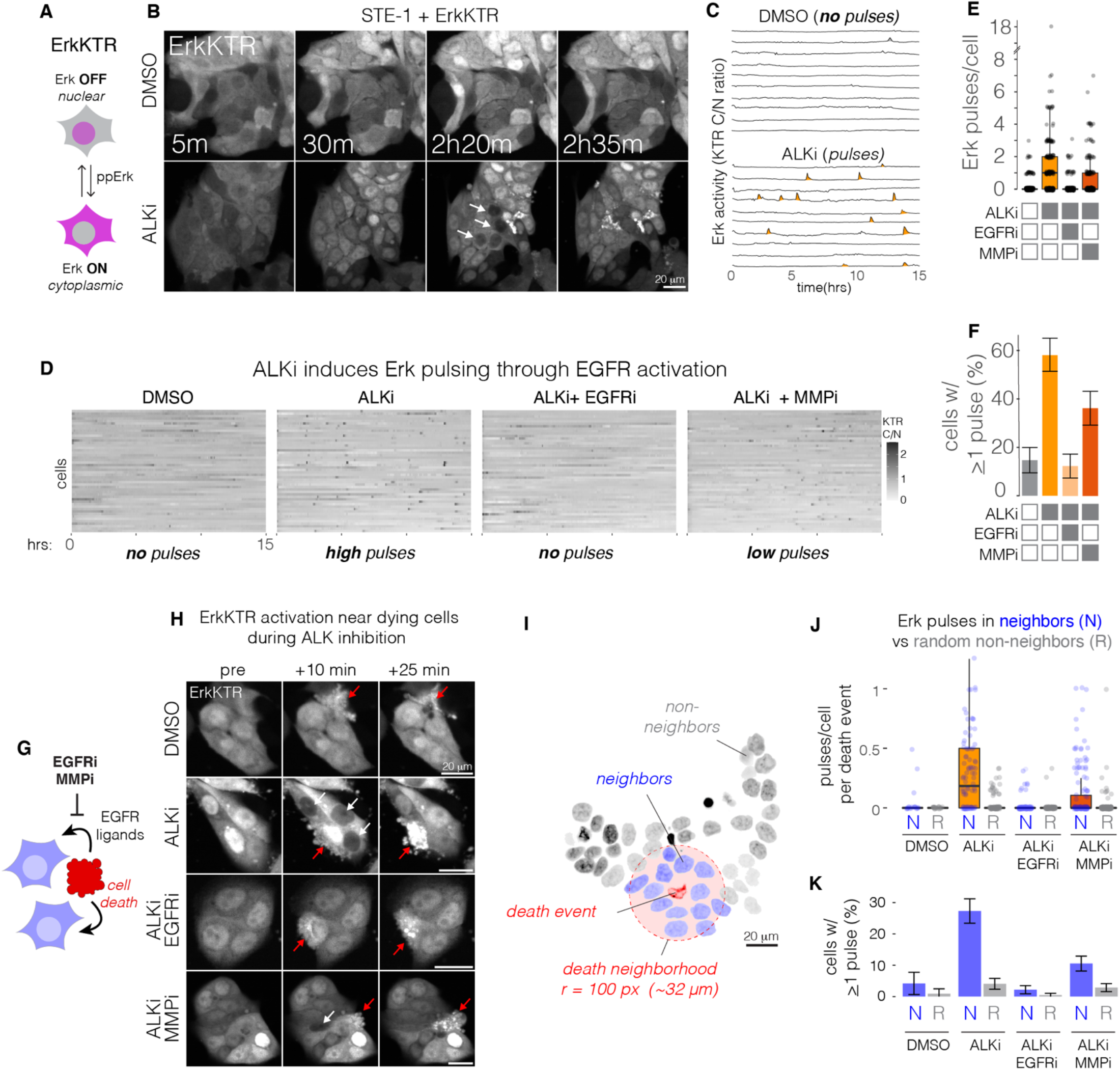
ALK inhibition hypersensitizes cancer cells to paracrine growth factors secreted from dying neighbor cells. A) The ErkKTR reporter indicates Erk activity through nuclear-cytoplasmic translocation of a fluorescent protein. B) Live cell imaging of STE-1 cells expressing ErkKTR in the presence or absence of ALKi (1 μM crizotinib). See **Supplementary Movie 3**. C) Representative single-cell traces of cytoplasmic/nuclear ErkKTR intensity ratio from conditions shown in B. D) Quantification of ErkKTR activity in the presence of ALKi or its combination with EGFRi (1 μM erlotinib) or MMPi (10 μM marimastat). E) Quantification of Erk activity pulses per cell. F) Fraction of cells that exhibited any pulses over 21.5 hrs of imaging. Error bars indicate 95% CI. n = 200 cells per condition. G) Apoptotic cells can secrete paracrine EGFR ligands to their neighbors. Paracrine signaling can be blocked by inhibiting either EGFR or the MMPs that are required to shed certain EGFR ligands from the surface of the sender cell. H) ErkKTR activity pulses are primarily observed surrounding a dying cell during ALK inhibition, but not in the absence of drug or in the presence of EGFR or MMP inhibitors. I) Definition of neighbors and non-neighbors of a death event. J) Quantification of pulses per cell for each death event in neighbors or a randomly chosen subset of cells not near a death event (see **Methods** for more details). K) Fraction of total neighbor vs random non-neighbor cells that show any pulsing. Error bars = 95% CI.

Activity pulses appeared sporadically but in a spatially coordinated manner, appearing either simultaneously or as a traveling wave within small clusters of neighboring cells (**Figure 5B, Supplementary Movie 3**). This pattern was consistent with RTK stimulation through paracrine signaling. To test this hypothesis, we sought to block paracrine signals through inhibition of either the EGF receptor (EGFRi, 1 μM erlotinib) or matrix metalloproteases (MMPi, 10 μM marimastat), which release EGFR ligands from the cell surface to enable paracrine signaling (Dong et al., 1999; Peschon et al., 1998; Sahin et al., 2004). Co-treatment with ALKi and EGFRi suppressed Erk pulses after drug addition (ALKi vs ALKi/EGFRi: 58 ± 7% vs 12 ± 5% of cells showing ≥ 1 pulse over 22 hrs; error = 95% CI), indicating EGFR activation causes the observed ERK pulses (**Fig. 5D,E,F, Supplementary Movie 3**). Co-treatment with ALKi and MMPi similarly reduced Erk reactivation pulses (ALKi vs ALKi/MMPi: 58 ± 7% vs 36 ± 7% of cells showing ≥ 1 pulse over 22 hrs; error = 95% CI), though to a lesser extent than with EGFRi, potentially due to MMP-independent juxtacrine signals (Brachmann et al., 1989; Wong et al., 1989) (**Fig. 5D,E,F, Supplementary Movie 3**). Thus, ALKi treatment in STE-1 cells decreases Erk signals but is rapidly followed by RTK reactivation mediated by paracrine signals.

### Signal reactivation results from paracrine signals from dying cells

We next sought to determine the source of the paracrine signals. We observed that pulsing events often appeared next to dying cells, and that co-inhibition of ALKi with EGFRi or MMPi prevented this pulsing (**Figure 5G,H, Supplementary Movie 3**). These observations are consistent with paracrine ligand secretion from apoptotic cells, which has previously been observed to promote cell survival and homeostasis within epithelial sheets (Aikin et al., 2020; Gagliardi et al., 2021; Valon et al., 2021). To quantify this effect, we measured signal activation in cells that neighbored a dying cell within the ~hour preceding its death (**Figure 5I**). We then counted Erk pulses in these neighbors (N) and compared pulse counts to those from randomly selected non-neighbors (R) over that same time interval (for more details, see **Methods**). Our analysis revealed that, in the absence of ALKi, pulses were almost never observed in either the N or R populations (DMSO, **Figure 5 J,K**). However, in ALKi-treated cells, N cells pulsed significantly more than R cells (N vs R: 27 ± 4% vs 4 ± 2% cells with ≥ 1 pulse; error = 95% CI) (**Fig. 5J,K**). Co-treatment with EGFRi eliminated Erk pulsing in neighbors (N cells in ALKi vs ALKi/EGFRi: 27 ± 4% vs 2 ± 1%), whereas co-treatment with MMPi dramatically reduced neighbor signaling relative to ALKi alone (N cells in ALKi vs ALKi/MMPi: 27 ± 4% vs 10 ± 2%),(**Fig. 5J,K**). Together, our results demonstrate that virtually all observed Erk reactivation is associated with paracrine signals associated with dying drug-treated cells. Importantly, because Erk pulses were not observed in untreated cells —even in neighbors of dying cells (**Figure 5J**)—these data suggest that ALKi-induced RTK hypersensitization is an essential first step for the perception of paracrine ligands during ALKi therapy.

### Signal reactivation pulses activate gene transcription

To understand the extent to which the short ALKi-dependent Erk pulses could impact cell behavior, we asked whether the observed pulses could stimulate downstream transcription. Erk activity is a strong driver of transcription, including of a class of rapidly responding immediate early genes (IEGs) that begin transcription within minutes of Erk activity(Wilson et al., 2017). EGR1 is an IEG that has been implicated in drug resistance to ALK inhibitors (Voena et al., 2013). Additionally, EGR1 expression is adaptive, such that its expression peaks by ~ 1 hr but then decays within 1-2 hours, even in the presence of constant upstream signal (Bugaj et al., 2018; Sukhatme et al., 1987) (**Figure S8A**). Thus, accumulation of EGR1 indicates the presence of only recent Erk activation(Davies et al., 2020). We thus examined the extent to which EGR1 accumulated in STE-1 cells upon drug treatment (**Fig. 6A**). In untreated cancer cells, EGR1 levels remained low despite high Erk signaling from EML4-ALK, consistent with EGR1 adaptation to tonic Erk signals (**Figure 6B,C**). Upon ALK inhibition, a distinct peak of EGR1-high cells appeared and grew at 4 and 6 hrs after drug treatment, indicating that ALKi-induced Erk pulses could indeed drive transcription. Importantly, co-inhibition of ALK and EGFR prevented the appearance of EGR1-positive cells, consistent with transcription resulting from paracrine signals through EGFR (**Figure 6B,C**). Thus, ALKi-induced Erk activity pulses provide sufficient signal to drive gene expression changes that could regulate cell fate.

**FIgure 6.**
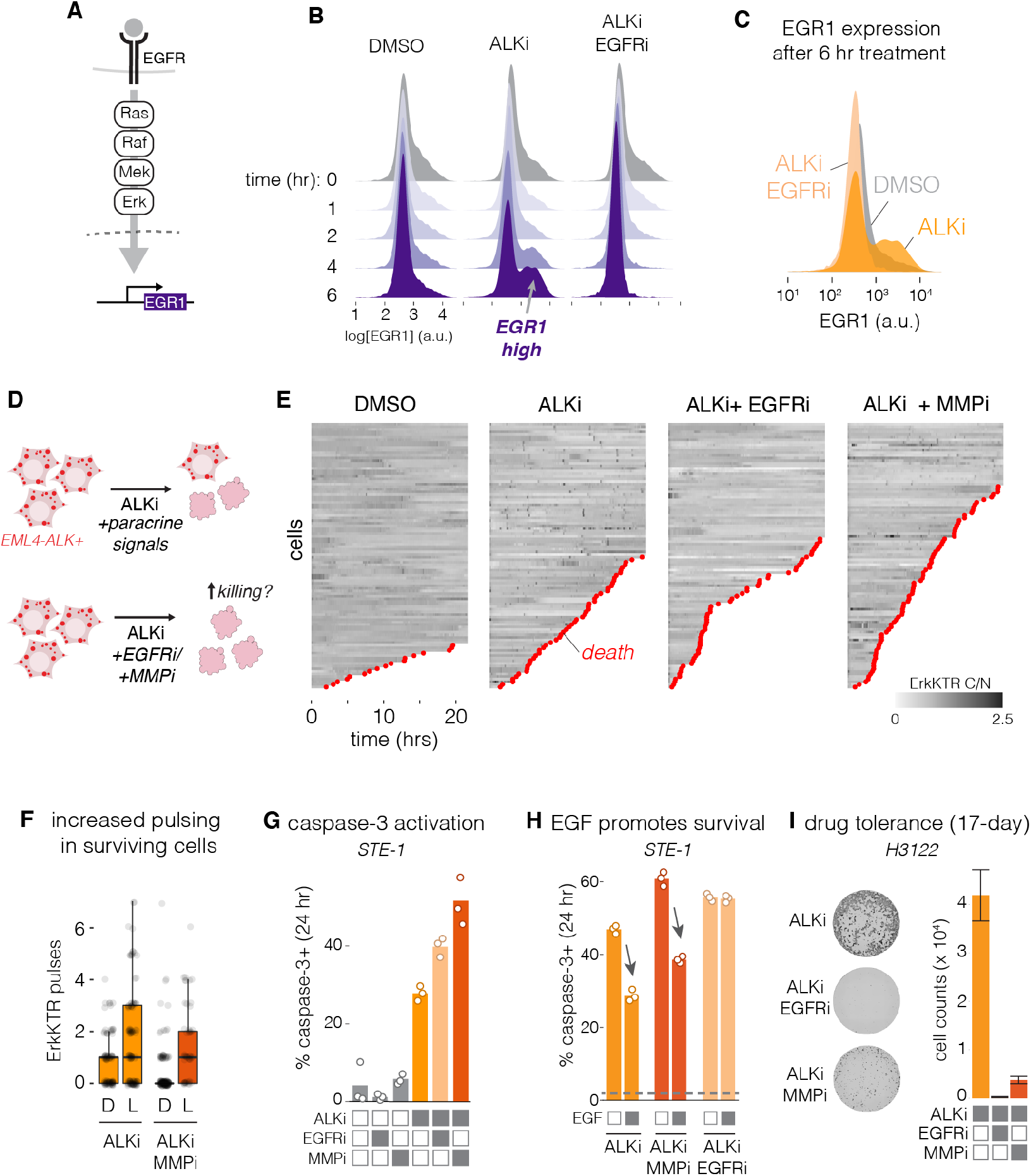
Resensitization to paracrine signals drives transcription and promotes drug tolerance. A) Signaling through EGFR activates Ras/Erk and stimulates transcription, including of the immediate early gene EGR1. B) Upon ALK inhibition, EGR1 expression increases, but not upon co-inhibition of ALK and EGFR. C) Comparison of EGR1 expression after 6 hrs of the indicated treatment. D) We hypothesized that restored perception of paracrine signals might promote survival, and that blocking paracrine signals would promote killing. E) Visualization of live cell imaging of Erk activity pulses and cell death. F) Quantification of pulses per cell in cells that died (D) or survived (L) through 22 hrs of imaging, in the conditions where Erk pulsing could be observed. G) Caspase-3 activation was assessed using the NucView reporter after 24 hr treatment with the indicated drugs. H) EGF (50 ng/mL) addition promotes survival during ALKi treatment and counteracts enhanced killing from ALKi/MMPi co-treatment, but not from ALKi/EGFRi co-treatment. Dotted line is cell killing from DMSO control treatment. Each data point represents the fraction of caspase-3+ cells from 2000-3000 cells. See **Methods** for more details. I) Crystal violet staining (left) and cell counts (right) of cell survival after 17 days of the indicated treatments.

#### Signal reactivation promotes acute drug tolerance and cell persistence during ALK inhibition

Finally, we asked whether RTK hypersensitization and resultant Erk pulses could counteract cell killing and promote drug tolerance to ALKi therapy (**Fig. 6D**). In STE-1 cells imaged over 22 hrs, ALKi treatment induced moderate cell death (45 ± 7% of cells; error = 95% CI). However, co-treatment of ALKi with either EGFRi or MMPi increased cell death (52 ± 7% and 75 ± 6% of cells, respectively) (**Fig. 6E**). In both ALKi- and ALKi/MMPi-treated cells, surviving cells showed an increased number of Erk pulses compared to cells that died (ALKi, live vs dying: 1.8 ± 0.2 vs 0.9 ± 0.1 pulses per cell; ALKi/MMPi, live vs dying: 1.6 ± 0.2 vs 0.4 ± 0.1 pulses per cell. Mean ± SE, **Figure 6F**), further drawing a link between RTK reactivation and cell survival. We confirmed our results through an independent assay of cell death using a fluorescent reporter of caspase-3 activity (NucView), an indicator of apoptosis, over the first 24 hr of treatment in EML4-ALK+ cells. As before, while ALK inhibition led to increased caspase-3, co-treatment with EGFRi significantly increased the caspase-3+ cell fraction in STE-1 cells (28 ± 2 vs. 41 ± 2%). Similarly, co-treatment with MMPi also increased caspase-3 (from 28 ± 2 to 50 ± 6%) (**Figure 6G**). Similar trends were observed in H3122 cells (**Figure S8B**). Neither EGFRi nor MMPi treatment alone showed enhanced killing over untreated cells. Notably, addition of 50 ng/mL EGF reversed enhanced killing in the ALKi/MMPi condition in STE-1 cells (61 ± 2 to 39 ± 1%, respectively), further suggesting that synergistic killing in the presence of MMPi resulted from blockade of signal reactivation (**Figure 6H**). Although EGF addition could not rescue enhanced killing under combined ALKi/EGFRi treatment, optogenetic pulses of optoFGFR lowered cell death in this condition (**Figure S8C,D,E**). Finally, we observed that co-treatments of ALKi with either EGFRi or MMPi suppressed long-term drug tolerance measured at 17-days (**Figure 6I**). Together, our results indicate that hypersensitization of EGFR and restored perception of paracrine ligands leads to reactivation of survival signals soon after drug treatment, which limits the cytotoxicity of ALK therapies and promotes drug tolerance, the first step towards acquired resistance. The mechanisms that underlie these events may thus provide novel treatment co-targets to enhance therapy in EML4-ALK+ cancers.

## Discussion

Our results advance a novel function for protein condensates in cancer cells and their response to targeted therapy (Jiang et al., 2020) (**Figure 7**). In addition to amplifying oncogenic signaling (Tulpule et al., 2021), EML4-ALK condensates simultaneously act as a molecular sponge for RTK adapters including Grb2 and SOS. This sequestration desensitizes the cell’s perception of external ligands by limiting the amount of adapters available to transduce the activated RTK signal. However, upon ALK inhibition, adapters are rapidly released to the cytoplasm, hypersensitizing cellular response to ligands in the cell’s microenvironment. RTK hypersensitization promotes cell survival and drug tolerance in response to targeted inhibitors, due at least in part to apoptosis-induced paracrine signals.

**Figure 7.**
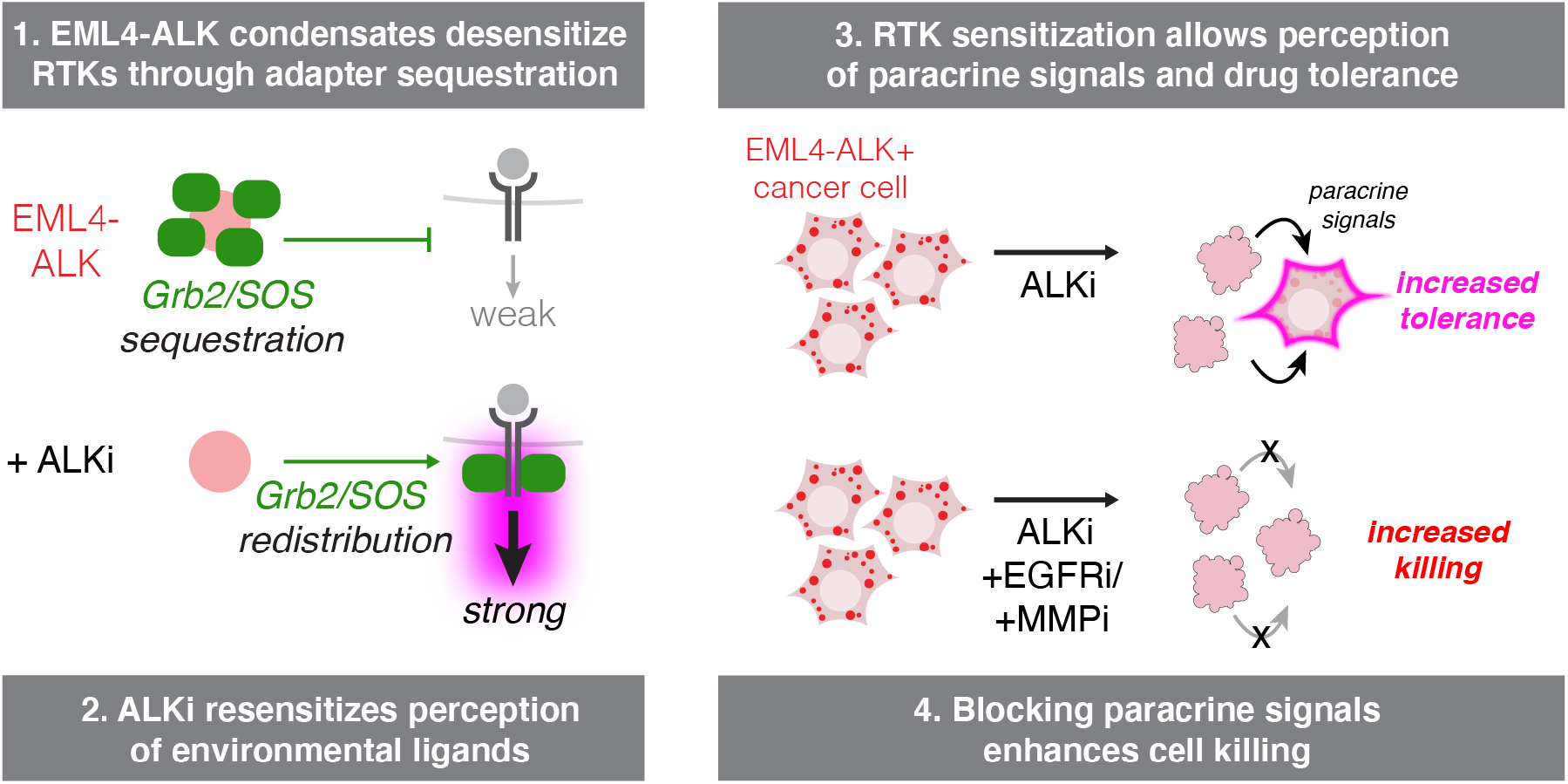
Model of how EML4-ALK condensates interact with transmembrane RTKs and its effect on drug tolerance. We find that EML4-ALK condensates suppress perception of EGFR signals by sequestering RTK adapters like Grb2 and SOS1. Upon ALK inhibition, adapters are released to the cytoplasm where they can transduce signals from EGFR and other RTKs. Although ALK inhibitors effectively suppress ALK activity, they simultaneously restore a cell’s perception to paracrine signals, including those from neighboring apoptotic cells. Such paracrine signals promote drug tolerance, and blocking them with inhibitors of either EGFR or MMPs enhances cell killing during ALK inhibitor therapy.

We show that protein condensation provides a mechanism by which oncogenes can suppress RTK signaling. Analogous RTK suppression has been observed in other, molecularly distinct cancers. BRAF V600E+ melanoma and colorectal cancer cells suppress RTKs through Erk-dependent transcription of negative regulators (Sproutys), and through inhibitory phosphorylation of SOS1(Gerosa et al., 2020; Lito et al., 2012; Prahallad et al., 2012). Inhibition of BRAF suppresses these mechanisms and leads to rapid, pulsatile Erk reactivation and drug resistance, similar to our observations. Separately, in cancers driven by EGFR, Erk provides suppressive phosphorylation of EGFR receptors, which is lost during MEK inhibition and leads to reactivation of ErbB3 and PI3K(Turke et al., 2012). In contrast to these and similar examples, our findings demonstrate that such RTK suppression can also be implemented through biophysical, rather than biochemical, feedback. Nevertheless, the variety of mechanisms by which oncogenes can suppress RTKs hints that such feedback might play an important role in establishing permissive conditions for oncogenesis. It will thus be interesting to more comprehensively understand the diverse oncogenic contexts in which such suppression and reactivation occurs.

It has been repeatedly observed that ligand-induced signaling through EGFR promotes survival and resistance to ALK inhibitors in EML4-ALK+ cancer cells (Obenauf et al., 2015; Sasaki et al., 2011; Tani et al., 2016; Tanimoto et al., 2014; Vaishnavi et al., 2017; Vander Velde et al., 2020; Wilson et al., 2012). Our work builds on these prior studies to show that 1) ALK inhibition is a necessary first step to allow cancer cell perception of paracrine signals, 2) EGFR ligands are sent from apoptotic neighbor cells, and 3) ALK-induced resensitization to RTK signals happens within minutes of drug treatment. In addition to fast RTK reactivation, previous work found that slower-timescale transcriptional amplification of KRas or downregulation of DUSP6 contributes to resistance development (Hrustanovic et al., 2015). Our results are complementary to these findings, which could be expected to further sensitize cells to external RTK ligands.

Dying cancer cells send survival signals to their neighbors through a mechanism that requires proteolytic processing of EGFR ligands, thus adding to previously identified mechanisms whereby drug treatment induces secretion that promotes cell survival and drug tolerance (Kurtova et al., 2015; Obenauf et al., 2015). Blockade of apoptosis-induced paracrine signals enhanced cell killing and limited tolerance in response to ALK inhibition. Although co-inhibition of EGFR is highly effective in promoting durable drug responses across a variety of cancer cells, (Sasaki et al., 2011; Vaishnavi et al., 2017; Vander Velde et al., 2020; Voena et al., 2013), a recent clinical trial of combined inhibition of crizotinib (ALKi) and erlotinib (EGFRi) in EML4-ALK+ NSCLC failed due to frequent adverse effects and low maximum tolerated dose (Ou et al., 2017). We show that co-inhibition of certain matrix metalloproteases may be an alternative strategy limit RTK signaling during targeted therapy. We note, however, that previous studies found that long-term MMP inhibition can promote resistance by causing accumulation of transmembrane RTKs (e.g. AXL) that are also MMP targets (Miller et al., 2016). Thus, further studies will be required to understand whether MMPi co-treatment can indeed enhance therapy, or whether MMPi scheduling could leverage acute benefits while avoiding deleterious chronic effects.

Our finding that the sequestration of RTK adapters can functionally desensitize RTK signals could inspire new types of therapeutic targets that mimic this behavior, for example through inhibition or sequestration of important RTK adapters. Of note, the recent discovery of EML4-ALK condensates has raised the idea that disaggregation of the condensates might be therapeutically beneficial (Cai et al., 2021; Hirai et al., 2020). Our work cautions, however, that such disaggregation strategies will likely be subject to the same adapter redistribution and rapid RTK reactivation observed with small molecule ALK inhibitors, potentially challenging their efficacy.

A unique promise of functional profiling is that common signaling abnormalities may be identified and inform therapies among genetically distinct cancers (Bugaj et al., 2017). Because our work revealed signaling principles that result from a likely common property (condensation) of RTK fusions (Du and Lovly, 2018; Tulpule et al., 2021), we anticipate that our findings may be broadly relevant to other cancer cells in this class. We show that in a cancer line driven by a CCDC6-RET fusion that also forms large aggregates (Tulpule et al., 2021), RET activity similarly suppresses EGFR, and RET inhibition potentiates EGFR signaling (**Figure S7**). These results are consistent with a previous study that showed that Grb2 can change association from the fusion to EGFR upon kinase inhibition in cancers driven by ALK, RET, NTRK1 and ROS1 fusions, including in patient samples of primary tumors, resistant tumors, and metastases(Vaishnavi et al., 2017). These findings imply common behavior across molecularly distinct RTK fusions and further point to the clinical relevance of these phenomena. Future work will determine the extent to which the principles we describe in EML4-ALK+ cancers will extend to this more diverse array of malignancies.

## Acknowledgements

The authors thank Dr. Magdalena Niewiadomska-Bugaj for feedback on statistics, data presentation and analysis. The authors also thank Dr. Alex Hughes (Penn), Dr. Sydney Shaffer, and Dr. Yael Mossé for helpful comments on the manuscript. This work was supported by the National Institutes of Health (R35GM138211 for L.J.B and D.G.M.; R01CA231300 for B.H. and T.G.B; U54CA224081, R01CA204302, R01CA211052 and R01CA169338 for T.G.B.), UCSF Marcus Program in Precision Medicine Innovation to B.H. and T.G.B, and the UCSF Physician Scientist Scholar Program and Alex’s Lemonade Stand for A.T. B.H. is a Chan Biohub Investigator.

## Author Contributions

D.G.M. and L.J.B. conceived the study. D.G.M. and L.R. performed experiments. J.G. and B.H. generated endogenously-tagged cell lines and provided guidance on imaging. D.G.M., T.R.M. and L.J.B. analyzed data. A.T., T.G.B., and L.J.B. provided experimental feedback and guidance. L.J.B. supervised the work. D.G.M and L.J.B. wrote the manuscript and made figures, with editing from all authors.

## Conflicts of interest

D.G.M. and L.J.B. have filed a provisional patent based on the findings in this work. T.G.B. is an advisor to Novartis, Astrazeneca, Revolution Medicines, Array/Pfizer, Springworks, Strategia, Relay, Jazz, Rain, Engine, Scorpion and receives research funding from Strategia, Kinnate, Verastem, and Revolution Medicines. A.T. is an advisor to Faze Medicines.

## Supplementary Figures

**Supplementary Figure 1.**
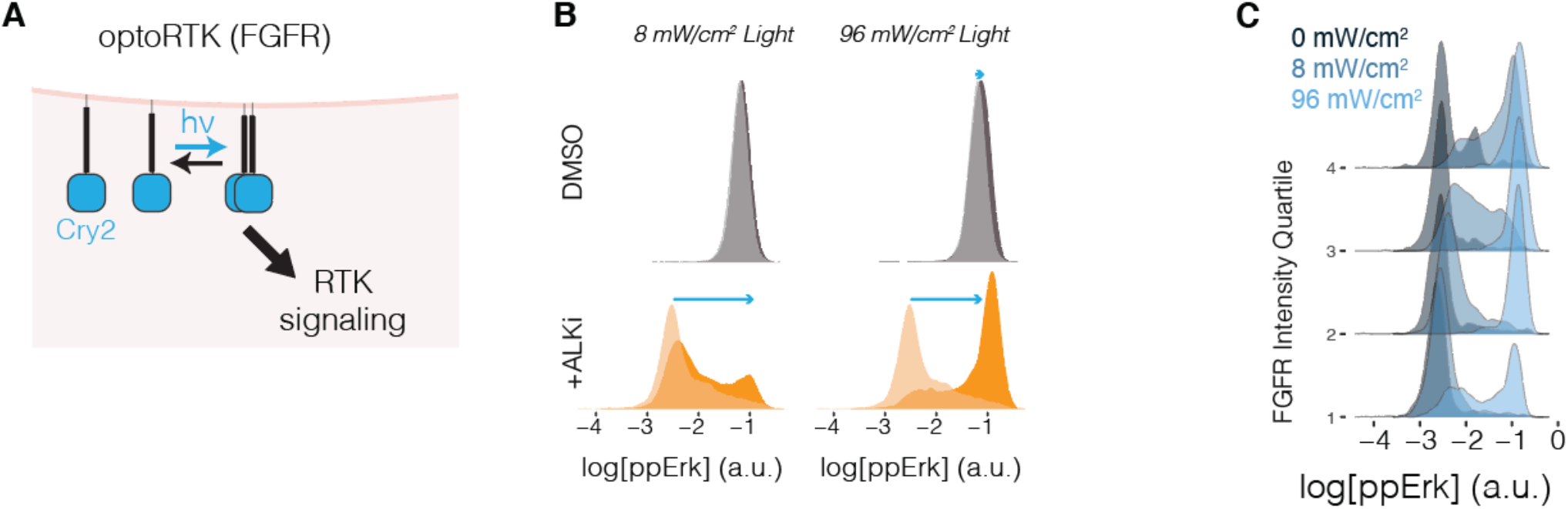
OptoFGFR functional profiling of STE-1 cancer cells. A) optoFGFR comprises the intracellular domain of FGFR1 fused to the PHR domain from Arabidopsis Cryptochrome 2, which clusters under blue light stimulation(Bugaj et al., 2013). The construct is anchored in the membrane through N-terminal myristoylation. B) Single cell distributions of ppErk intensity in STE-1 cells stimulated by the indicated intensity of blue light in the presence (orange) or absence (grey) of ALKi (2 hr pre-incubation of 1μM crizotinib). Strong ppErk response at low light intensity in the presence, but not the absence, of ALKi suggests that ALKi hypersensitizes cells to RTK stimulation. C) The magnitude of ppErk response in optoFGFR STE-1 cells is a function of both light intensity and optoFGFR expression levels. Results are presented by binning optoFGFR-mCh expression by quartiles. ( 1<25%, 25% < 2 < 50%, 50% <3 < 75%, 4 > 75%).

**Supplementary Figure 2.**
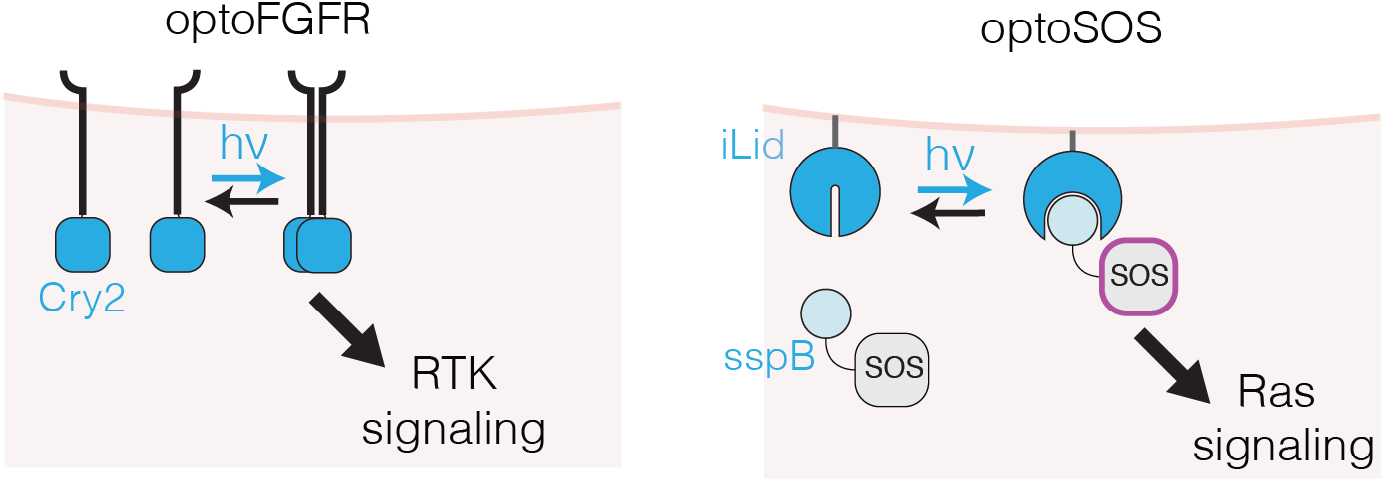
Schematics of the optoFGFR and optoSOS tools used in Figure 3. optoFGFR(Kim et al., 2014) allows optogenetic stimulation of the FGFR receptor using light-induced clustering of the Cryptochrome 2 protein(Bugaj et al., 2013). OptoSOS allos Ras activation through membrane recruitment of the SOS2 catalytic domain (SOS2_cat_). Membrane recruitment is achieved through blue light dimerization of sspB to the iLid protein(Guntas et al., 2015) which is anchored to the membrane through a CAAX motif.

**Supplementary Figure 3.**
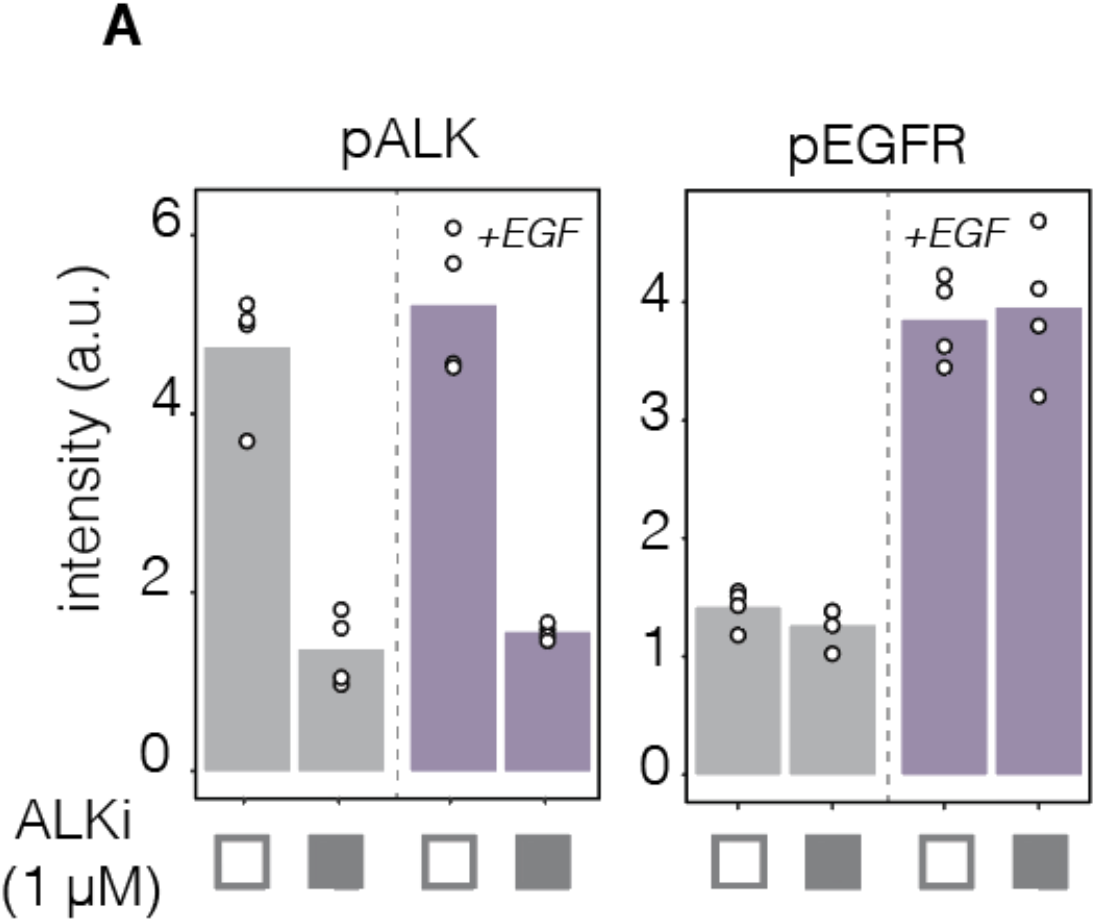
EGFR signal suppression does not result from decreased EGFR phosphorylation. Following overnight starvation, STE-1 cells were pretreated with ALKi or DMSO for 2 hrs. Cells were then stimulated with either EGF (50 ng/mL) for 15 min and then were fixed and immunostained for ppErk. ALKi pretreatment did not alter pEGFR (Y1068) levels in either the presence or absence of EGF, whereas pALK (Y1507) decreased. Thus EML4-ALK does not modulate EGFR signaling through changes in EGFR phsophorylation. Data points represent mean intensity of 3000-5000 cells.

**Supplementary Figure 4.**
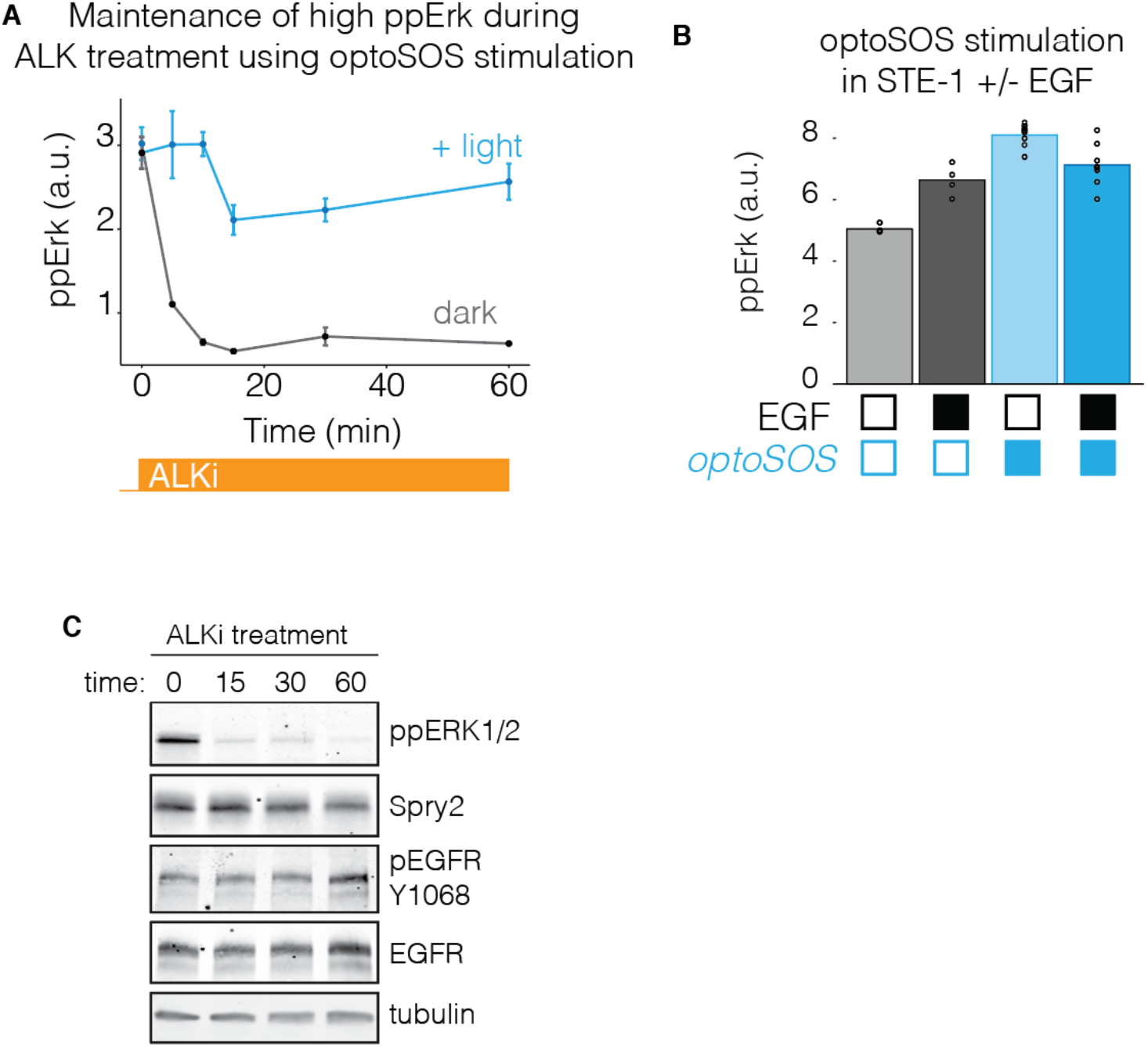
Determining the role of Erk-dependent negative feedback on RTK suppression. A) Demonstration that optoSOS can maintain high Erk levels (comparable to those from the uninhibited oncogene) during ALKi pre-treatment. Optogenetic stimulation of Ras/Erk was used to decouple ALK inhibition from loss of Erk signaling in **Figure 3G-J**. B) ppErk levels after 15 min of EGF stimulation in the presence or absence of optoSOS activation. Maintenance of optoSOS activation during EGF stimulation did not result in elevated Erk signaling in the absence of ALKi (as seen in **Figure 3J**). Data in A) and B) represent signal from top 25% of optoSOS-expressing cells. Data points represent mean ± SEM of 200-600 cells per condition. C) Western blot showing levels of negative regulator Spry2 over the first hour of ALKi treatment. Despite loss of Erk activity, Spry2 levels remain unchanged over this time period.

**Supplementary Figure 5.**
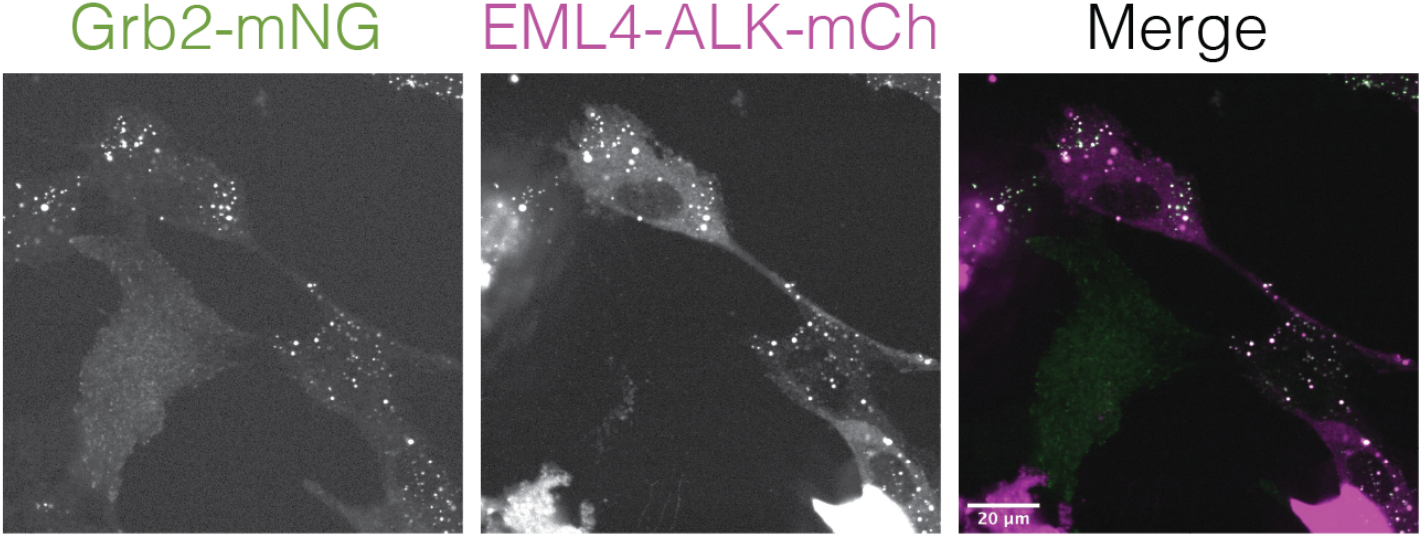
EML4-ALK condensates colocalize with Grb2 puncta. Beas2B cells that harbored endogenously tagged Grb2 (Grb2:mNG) were transfected with EML4-ALK-mCh. In transfected cells, EML4-ALK co-localized with Grb2. In untransfected cells, Grb2 was diffuse.

**Supplementary Figure 6.**
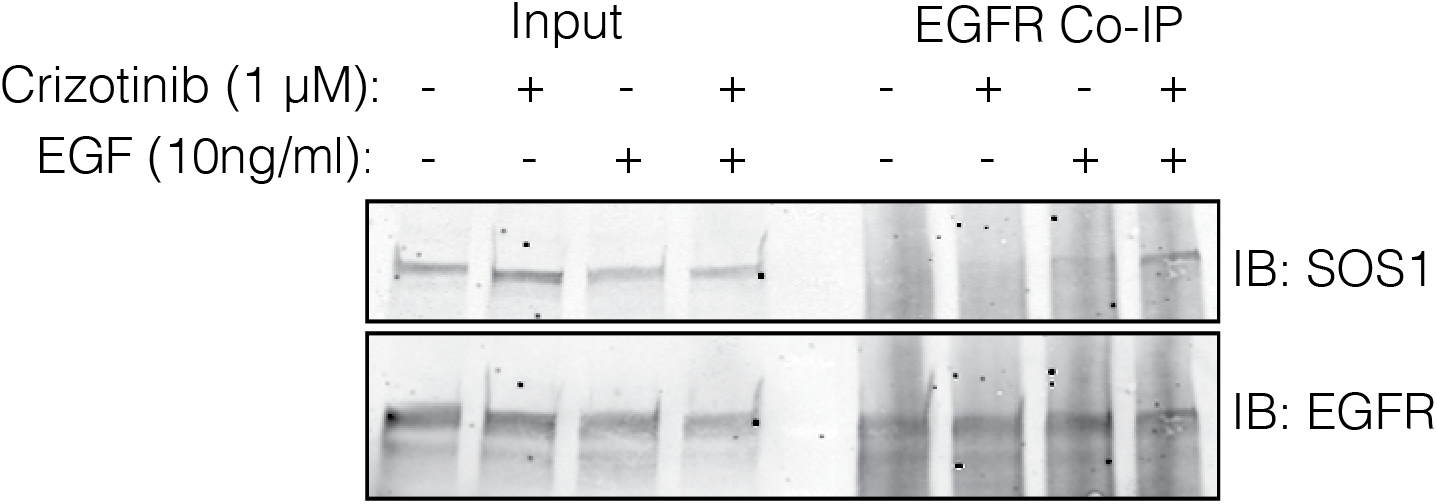
Co-immunoprecipitation of EGFR with SOS1. H3122 cells were starved overnight, pretreated with crizotinib (2 hr), and were stimulated with EGF for 2 minutes before lysis and immunoprecipitation. Right 4 lanes are reproduced from **Figure 4J**.

**Supplementary Figure 7.**
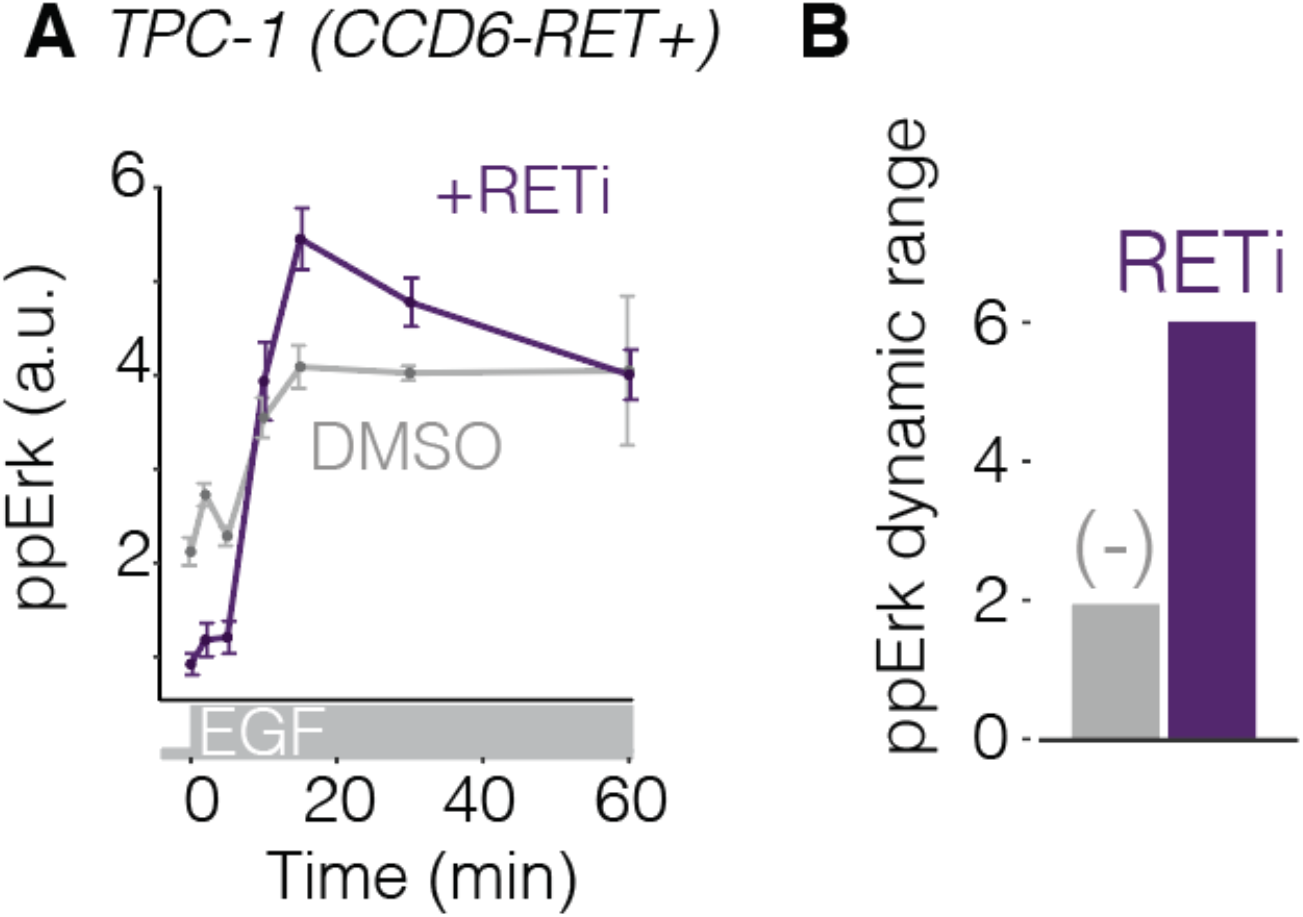
CCDC6-RET+ cancer cells show dynamic suppression of EGFR through RTK fusion activity. TPC-1 cells harbor CCDC6-RET, a distinct RTK fusion that forms condensates. We tested the extent to which CCDC6-RET could modulate EGFR dynamic range as observed for EML4-ALK. A) Plot shows ppErk response to EGF stimulation after 2 hr pre-treatment with DMSO (grey) or 100 nM BLU-667 (purple), a RET inhibitor. Data represent mean ± SEM of triplicates, each of which represents ~1000 cells. B) Quantification of ppErk dynamic range shows that oncogene inhibition enhances signaling through EGFR. Thus, multiple RTK fusions that form condensates dynamically suppress EGFR signaling.

**Supplementary Figure 8.**
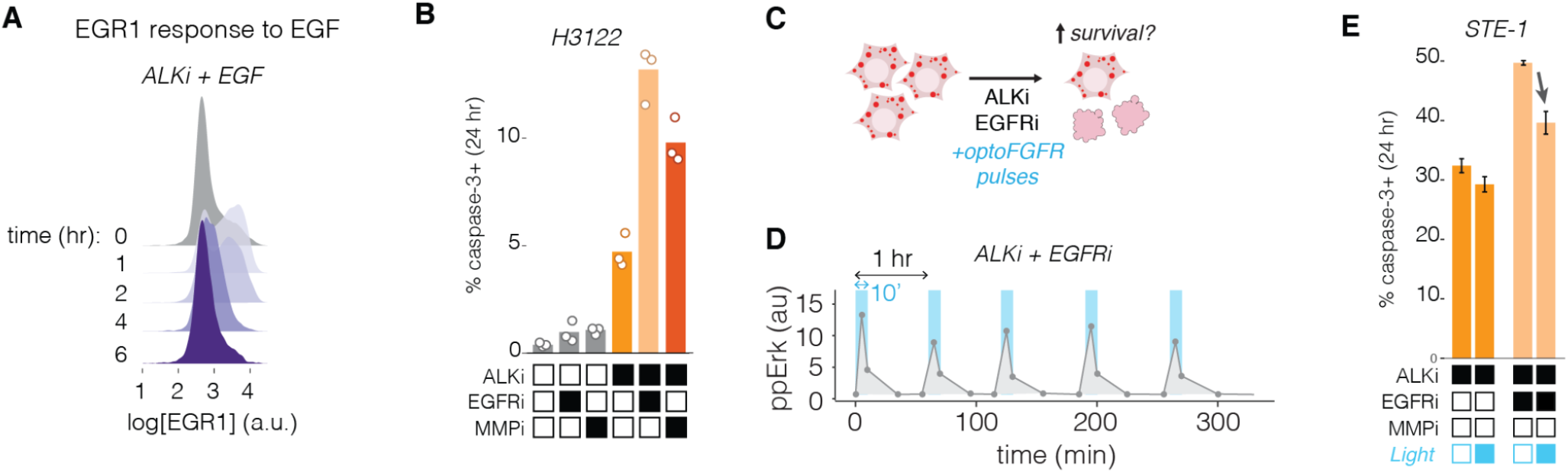
RTK/Erk pulses activate transcription and promote cell survival during ALKi treatment. A) Adaptive response of EGR1 expression in STE-1 cells treated with ALKi and 50 ng/mL of EGF. EGR1 peaks at 1 hr after stimulation and then adapts to near baseline expression. EGR response is stronger and earlier than observed in **Figure 6B** because here all cells were stimulated at t = 0, whereas in **6B** only individual cells were stimulated sporadically through paracrine signals starting 1-2 hrs after ALK treatment. B) Blockade of EGFR or MMPs accelerates cell death in H3122 in response to ALKi, as shown STE-1 cells in **Figure 6G**. C) We tested whether optoFGFR stimulation could reduce the enhanced cell death response in STE-1 cells co-treated with ALKi and EGFRi. D). A 10 min pulse of blue light (96 mW/cm^2^) every hour produced strong, periodic ppErk levels. Data points represent mean of three replicates, each representing ~2000 cells. E) Hourly pulses of optoFGFR/Erk (10 min of 96 mW/cm^2^) reduced cell death in both ALKi and ALKi/EGFRi treated cells. Data points represents the mean +/-SEM of three replicates, each representing ~10,000 cells.

## Methods

### Cell lines and cell culture

All cell lines were maintained at 37 °C and 5% CO_2_ using a standard cell culture incubator. STE-1, H3122, TPC-1, and Beas2B cells were cultured in RPMI-1640 growth medium (RPMI-1640 containing L-glutamine supplement with 10% fetal bovine serum (FBS) and 1% penicillin/streptomycin (P/S)). For experiments, cells were seeded in 96 or 384 well plates coated with fibronectin (MilliporeSigma, FC01010MG) diluted to 10 μg/mL in PBS. For 96 well plate experiments 10,000 STE-1 and H3122 cells were seeded in 200 μl of cell culture media per well. For 384 well plate experiments 5,000 STE-1, H3122, TPC-1 cells or 2500 Beas2B cells were seeded in 50 μl of culture media per well. LentiX-HEK 293T cells (TakaraBio, #632180) were cultured using DMEM supplemented with 10% FBS and 1% P/S.

### Plasmid design and assembly

All cloning was performed by PCR and DNA assembly using NEBuilder^®^ HiFi DNA Assembly Master Mix (New England Biolabs #E2621). Constructs for transient expression were cloned into pEGFP-C1 backbones encoding a CMV promoter. Fluorescently-tagged EML4-ALK (pCMV EML4-ALK-GFP) was generated by inserting the coding sequence for EML4-ALK(V1) upstream of GFP. To generate untagged EML4-ALK for transient expression in Beas2B cells, EGFP was replaced with P2A-H2B-iRFP (pCMV EML4-ALK-P2A-H2B-iRFP). EML4-ALK (ΔTD) mutant was cloned by excluding bases 310-459 from pCMV-EML4-ALK-P2A-H2B-iRFP through PCR amplification followed by plasmid assembly using HiFi. EML4-ALK (K589M) was cloned by single point mutation encoded on the primer used to amplify pCMV-EML4-ALK-P2A-H2B-iRFP followed by blunt end ligation using NEB T4 ligase (NEB, #M0202). Constructs for generation of stable cell lines were cloned into pHR lentiviral or CLPIT retroviral plasmid backbones. For optogenetic control of FGFR signaling (Kim et al., 2014), the FGFR intracellular domain with a N-terminal myristoylation site was inserted upstream of mCh-Cry2(Bugaj et al., 2013) in a CLPIT plasmid to create CLPIT Myr-mCh-FGFR(ICD)-Cry2. For optogenetic control of SOS signaling, we generated pHR sspB-SOScat-mCh-2A-iLid-CAAX by inserting an mCherry coding sequence to replace BFP, which we described previously (Benman et al., 2022). For live-cell tracking of ppErk activity, we generated pHR ErkKTR-mRuby2 by inserting an mRuby2 coding sequence in place of BFP in a construct we described previously (Benman et al., 2022). Visualization of nuclei was achieved using pLenti PGK DEST-H2B-iRFP670 (Addgene plasmid #90237).

### Transient transfection

pCMV-EML4-ALK-P2A-H2B-iRFP, pCMV-EML4-ALK(ΔTD)-P2A-H2B-iRFP, pCMV-EML4-ALK(K589M)-P2A-H2B-iRFP and CMV-3XFLAG-mCh-EML4-ALK were transiently transfected into wt Beas2B or Beas2B mNG:Grb2 cells using Lipofectamine™ 3000 (Invitrogen, L3000001). Briefly, 2.5×10^3^ cells were seeded in fibronectin-coated 384 well plates in RPMI growth medium. For an individual well: 0.25 μl of Lipofectamine 3000 reagent was diluted in 1 μl of OptiMEM (Gibco, #31985070) and incubated at room temperature for 5 min. This mix was then added to a diluted DNA mix composed of 25 ng of plasmid DNA and 0.5 μl of P3000 reagent, diluted in OptiMEM to a final volume of 1.25μl. After 15 min of incubation at room temperature, transfection mix was added to cells cultured in 50 μl culture medium. Following 6 hrs of transfection, media was exchanged with fresh growth medium. Cells were then starved for at least 6 hrs using starvation medium (RPMI + 1% P/S) with or without phenol red for fixed or live-cell assays, respectively.

### Generation of stable cell lines

Cell lines were generated using viral transduction. CLPIT-Myr-mCh-FGFR(ICD)-Cry2, pHR sspB(nano)-mCh-SOS2(cat)-2A-iLid-CAAX, pHR KTR-mRuby2, and PGK-DEST-H2B-iRFP670 vectors were packaged into lentivirus using HEK 293T cells. Briefly, 7.0 × 10^5^ cells were seeded in each well of a 6 well plate. For pHR transfer vectors, cells were transfected with 1.5 μg of pHR transfer vector, 1.33 μg of pCMV-dR8.91 (Addgene #12263), and 0.17 μg pMD2.G (Addgene #12259). For CLPIT transfer vectors, cells were transfected with 1.25 μg of transfer vector, 0.5 μg of pCMV-VSVG (Addgene #8454) and 0.75 μg of pCMV-gag/pol. Transfections were performed using the calcium phosphate method. The following day, medium was removed and replaced with fresh growth medium. After 24 and 48 hrs, virus-containing supernatant was collected and stored at 4 °C. Supernatants were then centrifuged at 200 x g for 2 min to remove cell debris, filtered through a 0.45 μm filter (Fisher Scientific, catalog number 13-1001-07), and used immediately or stored at −80°C. For transduction, 2.0 × 10^5^ H3122 or STE-I cells were seeded along with 500 μl of viral supernatant. Transduced cells were expanded and sorted (BD Influx) for appropriate expression levels. For STE-1 cells that expressed optoFGFR or optoSOS, the top 20% or 30% of mCh+ cells, respectively were sorted, expanded, and used for experiments. Beas2B cells expressing endogenously tagged Grb2:mNG (Beas2B Grb2:mNG) generated previously (Tulpule et al., 2021).

### Optogenetic Stimulation

Light stimulation was achieved in microwell plates using the optoPlate-96 (1-color blue version(Bugaj and Lim, 2019)). A 20 mm tall black adaptor was used to ensure even light diffusion across each of the 384 well plate wells. To investigate the ppErk response to optoFGFR stimulation (**Figure 1 F,G**), cells were stimulated with 500 ms every 10 sec for 5 min (light intensity ranging from 3.2mW/cm^2^ (min) to 160mW/cm^2^ (max)). To maintain basal ppErk levels during ALK inhibition (**Figure 2 H,J**) optoSOS-expressing STE-1 cells were stimulated for 500ms every 10 sec (160 W/cm^2^ blue light) for 15 min, with stimulation beginning simultaneously with drug treatment (1μM crizotinib). Cells were then stimulated with 50 ng/ml EGF (PeproTech, 315-09) while still being exposed with blue light as previously described. For optogenetic rescue of cell death during ALKi/ EGFRi treatment, cells were stimulated with blue light (96 mW/cm^2^, 500msON/10s OFF) for 10 min every hour, with stimulation beginning simultaneously with drug treatment.

### Growth Factor Stimulation Assay

Plated cells were starved overnight with RPMI-1640 starvation medium. Starvation was achieved by performing 7X 80% washes either manually or using the BioTek 405 LS microplate washer. Cells were treated with inhibitor for 2 hours (1μM crizotinib (Sigma-Aldrich, PZ0191) or 0.1μM BLU-667 (Biovision, B2548)). Cells were then stimulated at their respective time points with EGF following a delay that allowed for all wells to be fixed simultaneously. Cells were fixed with 4% paraformaldehyde (PFA) (Electron Microscopy Sciences, 15710) by adding the appropriate volume of 16% PFA to the culture medium to yield a final concentration of 4%. Samples were then immediately prepared for immunostaining.

### Immunofluorescence

After PFA fixation (10 mins), cells were permeabilized with 0.5% Triton-X100 in PBS for 10 min, followed by incubation in ice-cold 100% methanol for 10 min. Samples were then blocked with blocking solution (1% bovine serum albumin (BSA) (Fisher, BP9706100) diluted in PBS) for 1 hour at room temperature. Samples were incubated in primary antibody diluted in blocking solution for either 2 hours at room temperature (RT) or overnight at 4 °C. Primary antibodies used were: phospho-p44/42 MAPK (Erk1/2) (Thr202/Tyr204), Cell Signaling #4370, 1:400 dilution; phospho-EGF Receptor (Tyr1068), Cell Signaling #3777, 1:800; EGR1, Cell Signaling # 4153, 1:800). After incubation with primary antibody, samples were washed 7X with 80% washes of 0.1% Tween-20 in PBS (PBS-T) using the BioTek 405 LS microplate washer. Samples were then incubated in blocking solution containing secondary antibody (IgG (H+L) Cross-Adsorbed Goat anti-Rabbit, DyLight™ 488, Invitrogen #35553, 1:500; Goat anti-Rabbit IgG (H+L) Cross-Adsorbed Secondary Antibody, DyLight™ 650, Invitrogen #SA510034, 1:500) and 4,6-diamidino-2-phenylindole (DAPI; ThermoFisher Scientific #D1306, 300 nM) for 1 hour at RT. Samples washed with PBS-T as previously described. Samples were left in a final volume of 100 μL PBS-T for imaging and storage.

### Immunoblotting and co-Immunoprecipitation

For immunoblots STE1 or H3122 cells (3 × 10^5^) were plated in each well of a 6 well plate, cultured for 24 hours, and subsequently serum starved for 16 hours. Cells were treated with 1 μM crizotinib for the times indicated, after which cells were washed in ice cold PBS and lysed in RIPA buffer (50 mM Tris pH 7.5, 150 mM NaCl, 1% NP40, 0.1% SDS, 0.5% DOC, 1 mM EDTA, 2 mM sodium vanadate and protease inhibitor). Protein concentration of cleared cell lysates was determined by BCA protein assay kit (Thermo Scientific #23225) and 20-30 μg of lysed samples were subjected to SDS-polyacrylamide gel electrophoresis (SDS-PAGE). For co-immunoprecipitation, STE1 or H3122 cells (2.5 × 10^6^) were plated on 10 cm plates, cultured for 24 hours, and subsequently serum-starved for 16 hours. Cells were treated with 1 μM crizotinib and 10 ng/mL EGF as indicated, washed with ice cold PBS, and lysed (50 mM HEPES pH7.4, 150mM NaCl, 1% Triton X-100, 1 mM EDTA, 1 mM EGTA, 10% glycerol, 2 mM sodium vanadate and protease inhibitor (Sigma #P8340)). Cleared cell lysates were incubated for 2 hours with Protein A/G agarose beads (Santa Cruz, SC-2003) that were hybridized with EGFR antibody (Thermo, clone H11). Beads were then washed 5 times with HNTG buffer (20 mM HEPES pH 7.4, 150 mM NaCl, 0.1% Triton X-100, 10% glycerol), and sample buffer was added to elute proteins. Eluates or 25-30 μg of protein lysate were loaded in a precast 4-15% gradient SDS-polyacrylamide gel for electrophoresis (mini-protean TGX precast gel, Bio-RAD, # 456-1084).

Protein separations were transferred onto a nitrocellulose membrane using the Trans-blot Turbo RTA transfer kit (Bio-rad, #170-4270) according to manufacturer’s protocol. Membranes were blocked in 5% milk in Tris buffer saline with 0.5% Tween-20 (TBS-T) for 1 hour and incubated overnight at 4°C with primary antibodies against EGFR (CST #4267), SOS1 (CST #5890), SPRY2 (CST #14954), pERK1/2 (CST #4370), tubulin (CST #3873). Each primary antibody was used at a dilution of 1:1000 in TBS-T with 3% BSA. After washing with TBS-T, membranes with incubated with secondary antibodies in TBS-T with 3% BSA for 1 hr at room temperature (IRDye^®^ 800CW Goat anti-Rabbit IgG, 1;20,000 dilution, LI-COR #926-32211; IRDye^®^ 680RD Donkey anti-Mouse IgG, 1:20,000 dilution, LI-COR, #926-68072). Membranes were then imaged on the LI-COR Odyssey scanner.

### Live cell imaging

Live cell imaging was performed using a Nikon Ti2-E microscope equipped with a Yokagawa CSU-W1 spinning disk, 405/488/561/640 nm laser lines, an sCMOS camera (Photometrics), and a motorized stage. Cells were maintained at 37 °C and 5% CO_2_ using an environmental chamber (Okolabs). Transfected Beas2B Grb2:mNG cells were imaged 48 hours after transfection with a 40X oil immersion objective. To visualize Grb2 localization during drug response and EGF stimulation, cells were imaged for 2 hours following addition of 1 μM crizotinib and subsequent EGF stimulation, both added manually during image acquisition. For monitoring Erk activity after drug treatment, STE-1 cells stably expressing ErkKTR-mRuby2 and H2B-iRFP670 were seeded in a 96 well plate (Falcon #353072) at a density of 1.5×10^4^ cells/well. The following day, media was replaced with phenol-free RPMI-1640 with L-glutamine supplemented with 2% FBS and 1% P/S. Immediately before imaging, cells were supplemented with NucView (Biotium #10402, 1 μM) and the indicated drugs (crizotinib (1 μM), erlotinib (1 μM, Sigma-Aldrich, #SML2156), marimastat (10 μM, Cayman, #14869)) or DMSO (1:5000)). Cells were imaged every 5 minutes for 22 hours using confocal microscopy using 20X magnification.

### Caspase-3 Activation and Drug Tolerance Assay

To assay acute cell killing, STE-I and H3122 cells were seeded in a 96 well plate. The following day, media was replaced with phenol-free RPMI-1640 supplemented with 2% FBS, 1% P/S, and treated with 1μM NucView 488 and the indicated (1μM crizotinib, 1μM erlotinib, 10μM marimastat). 24 hours after drug addition, cells were incubated in 5 μg/mL Hoechst 33342 Hydrochloride (Cayman 15547) for 15 minutes. Cells were then imaged under live-cell confocal microscopy using 10X magnification. To assay drug tolerance, H3122 cells were seeded in a 96 well plate as previously described. The following day, media was replaced with RPMI with 2% FBS, 1% P/S and treated with the indicated inhibitors (1μM crizotinib, 1μM erlotinib, 10μM marimastat). Media with inhibitors was replaced every 2 days. Following 17 days of treatment, cells were fixed with 4% PFA and permeabilized with 100% methanol for 10 min. Cells were then incubated in DAPI (300 nM) for 30 min, imaged, and nuclei were counted through image analysis

### Image processing and analysis

#### Immunostaining and caspase-3 reporter

Images of fixed-cell immunostaining and live-cell caspase-3 reporter were quantified using CellProfiler (v 4.0.7)(Lamprecht et al., 2007). Briefly, cell nuclei were segmented using the DAPI channel, and the cytoplasmic fluorescence was measured within a 5-pixel ring that circumscribed the nucleus. For measuring caspase-3 activity and drug tolerance, DAPI or Hoechst-stained nuclei were segmented using ilastik(Berg et al., 2019), quantified in CellProfiler, and quantification was exported to R for processing and data visualization using the tidyR package (Wickham et al., 2019).

#### Live cell imaging of Grb2

Time lapse imaging of Grb2 puncta was quantified with a custom MATLAB script. Briefly, cells were segmented manually and clusters were identified in a semi-automated manner through the following steps. First, cell intensities were normalized to their internal median. Images were then passed through sequential Gaussian, top hat, and Laplacian filters to enhance clusters while suppressing other features like background and high frequency noise. These transformed intensities were used to identify bright pixels at the center of clusters with a user-defined intensity cutoff. From each of these center points, neighboring pixels were compared to a threshold intensity that was set by the local background of the cell and scaled by a second user-defined parameter. Neighboring pixels above this threshold are included in the cluster, while pixels below the threshold are excluded. This occurs iteratively, where adjacent pixels to each included pixel were also checked to be included or excluded until all new neighbors were below the threshold. Cluster properties of each cell were exported and processed in R for analysis and visualization.

#### Live cell imaging of ErkKTR

ErkKTR dynamics and cell death were tracked and quantitated in a semi-automated manner using the p53CellCinema package (Reyes et al., 2018) in MATLAB, and data was processed and visualized in R and RStudio using the tidyR package. Briefly, cells with low or no ErkKTR expression were first excluded from analysis. Then, ErkKTR cytoplasmic/nuclear ratios from remaining cells were imported into MATLAB, and peaks were identified using the ‘findpeaks’ function. Peak calls were then manually inspected for outliers, for example resulting from apoptotic cell debris, and outlier cells were removed from analysis. Of note, we consistently observed Erk pulses preceding cell division events across conditions. This phenomenon has been previously observed and has been shown to be independent of Ras/Erk signaling, potentially due to non-specific activation from a cyclin dependent kinase(Gerosa et al., 2020). Thus, we disregarded all pulses that occurred within 10 frames (50 mins) preceding cell division. To count Erk pulses in neighbors vs non-neighbors of dying cells, we first identified each cell death event. We then identified the neighbors of the dying cell by identifying which nuclear centroids were within a 100 pixel radius (~32 μm, or 2-3 cell diameters) of the dying cell for each of the 10 frames preceding the death event. We then counted the total pulses of those neighbors over the indicated time frame, and compared against pulse counts from a random subset of non-neighbor cells over the same time frame. Importantly, for each death event, the number of non-neighbors matched the number of neighbors, and the sampled non-neighbors were not neighbors of other death events over that same time span. For visualization in **Figure 6E**, cells that divided during the course of the experiment are represented as separate cells, and the Erk history of the mother cell is reproduced for both cells. However, for quantification of Erk pulses, mother cell pulses were counted only once.

